# Characterisation of a TatA/TatB binding site on the TatC component of the *Escherichia coli* twin arginine translocase

**DOI:** 10.1101/2022.12.12.520026

**Authors:** Emmanuele Severi, Mariana Bunoro Batista, Adelie Lannoy, Phillip J. Stansfeld, Tracy Palmer

## Abstract

The twin arginine transport (Tat) pathway exports folded proteins across the cytoplasmic membranes of prokaryotes and the thylakoid membranes of chloroplasts. In *Escherichia coli* and other Gram-negative bacteria, the Tat machinery comprises TatA, TatB and TatC components. A Tat receptor complex, formed from all three proteins, binds Tat substrates, which triggers receptor organisation and recruitment of further TatA molecules to form the active Tat translocon. The polytopic membrane protein TatC forms the core of the Tat receptor and harbours two binding sites for the sequence-related TatA and TatB proteins. A ‘polar’ cluster binding site, formed by TatC transmembrane helices (TMH) 5 and 6 is occupied by TatB in the resting receptor and exchanges for TatA during receptor activation. The second binding site, lying further along TMH6 is occupied by TatA in the resting state, but its functional relevance is unclear. Here we have probed the role of this second binding site through a programme of random and targeted mutagenesis. Characterisation of three stably produced TatC variants, P221R, M222R and L225P, each of which is inactive for protein transport, demonstrated that the substitutions did not affect assembly of the Tat receptor. Moreover, the substitutions that we analysed did not abolish TatA or TatB binding to either binding site. Using targeted mutagenesis we introduced bulky substitutions into the TatA binding site. Molecular dynamics simulations and crosslinking analysis indicated that TatA binding at this site was substantially reduced by these amino acid changes, however TatC retrained function. While it is not clear whether TatA binding at the TMH6 site is essential for Tat activity, the isolation of inactivating substitutions indicate that this region of the protein has a critical function.

## INTRODUCTION

The general secretory (Sec) and twin arginine translocase (Tat) systems operate in parallel to export proteins across the cytoplasmic membranes of prokaryotes and the thylakoid membranes of plant chloroplasts (1, 2). While the Sec pathway transports unfolded proteins, substrates of the Tat pathway are exported in a folded state (3, 4). Proteins are targeted to the Sec or Tat machineries by the presence of a signal peptide at their N-terminus, which is usually cleaved during transport (5). Sec and Tat signal peptides are superficially similar, but each has features that ensure passenger proteins are targeted to the appropriate transport system (6–8). In particular Tat signal peptides contain an almost invariant twin arginine motif that is critical for recognition by the Tat machinery (7, 9).

In prokaryotes, the Tat system is best studied in the model bacterium *Escherichia coli*. The *E. coli* Tat translocase comprises three membrane proteins, TatA, TatB and TatC (10–13). TatE, which is produced at very low levels is a minor component that is functionally equivalent to TatA and dispensable for Tat activity in laboratory conditions (10, 14). Although TatA and TatB have differing functions during Tat transport, they derive from the same protein family (15). They are monotopic membrane proteins with a very short N-terminal transmembrane helix, followed by an amphipathic helix located at the cytoplasmic side of the membrane, and an unstructured C-terminal tail (16, 17). TatB proteins are generally bigger than TatA, having a longer amphipathic helical region and more extended tail (15) (Fig 1A). TatC is the largest Tat component, consisting of six transmembrane domains with the N- and C-termini in the cytoplasm (9, 18) (Fig 1A).

**Fig 1.**
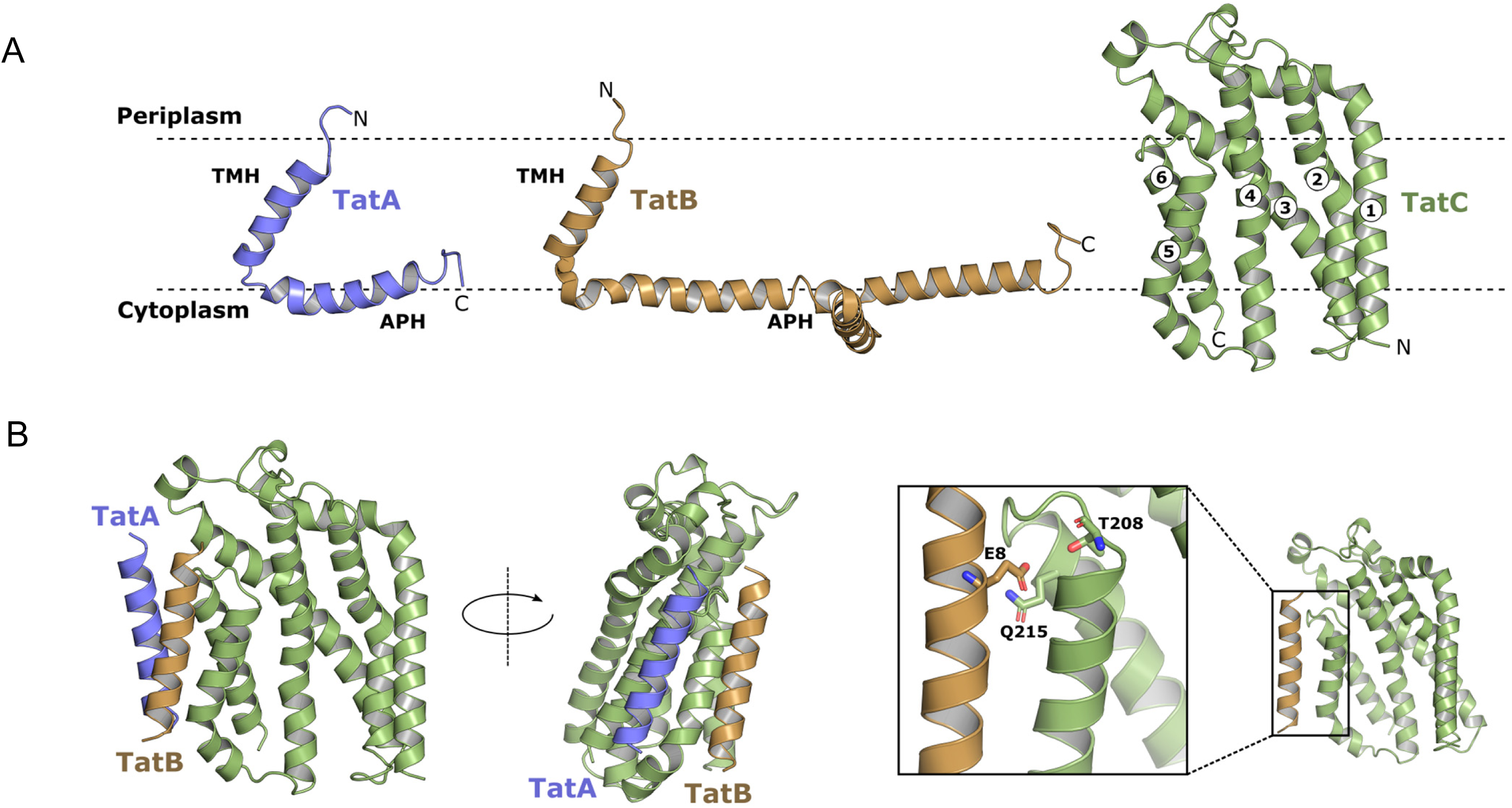
(a) Structures of the TatA (blue), TatB (orange) and TatC (green). The transmembrane helices (TMHs) and amphipathic helices (APHs) of TatA and TatB are indicated, together with the TMH numbering in TatC. (b) A model for the resting state of the TatABC receptor complex, showing the interactions of the transmembrane helices, with the constituent subunits coloured as in part A. (c) Polar cluster interactions between TatC residues T208 and Q215 (green) and TatB residue E8 (orange).

The Tat system is dynamic, and the active translocase assembles ‘on demand’, in a proton-motive force-dependent manner, triggered by functional interaction with substrate proteins (19–21). In the resting state, the machinery comprises a TatABC ‘receptor complex’ with many further TatA molecules dispersed throughout the membrane (21–23). The Tat receptor complex is multimeric, containing several copies each of Tat component, likely present in a 1:1:1 ratio (15, 22, 24). Crosslinking, mutagenesis and co-evolution analysis has identified a critical binding site for TatA/TatB at transmembrane helix (TMH) 5 of TatC (15, 25, 26). Binding to this site is mediated by a polar cluster of residues located at the C-terminal end of TatC TMH5 through to the start of TMH6 which co-ordinate a polar residue present, at equivalent positions, in the TMH of TatA or TatB. While either protein is able to bind to occupy this binding site, in the resting state it is occupied by TatB (15)(Fig 1B). Assembly of the translocase is initiated by interaction of the Tat receptor with a signal peptide. The conserved twin arginine motif is recognised by a negatively charged surface patch on TatC (9). The signal peptide subsequently binds more deeply within the receptor, making extensive contacts with the TatB TMH (20, 25, 27–30). This is accompanied by a rearrangement at the TMH5 binding site, where TatA replaces TatB, priming the recruitment of additional TatA protomers from the membrane pool (19–21, 31–34). Substrate transport across the membrane is facilitated by the TatA oligomer, potentially through localised weakening of the cytoplasmic membrane (9, 16, 35).

Unexpectedly, a second binding site for TatA and TatB was also identified on TatC TMH6, through crosslinking analysis (26). Again, each protein was able to bind to this site, but TatA was shown to occupy the site under resting conditions (26). The functional relevance of this second binding site is currently unclear, because very few TatC-inactivating mutations have been identified that fall in this region of the protein. To explore the role of this binding site further we have undertaken extensive mutagenesis of TatC TMH6 and biochemical analysis of variant Tat receptor complexes. Our results indicate that this site is surprisingly robust to amino acid substitution and that even multiple bulky substitutions in positions that would be expected to disrupt binding interfaces do not abolish Tat activity.

## METHODS

### Strains and Plasmids

*E. coli* strain MC4100 (36) and the isogenic *tat* deletion mutants DADE (as MC4100, Δ*tatABCD*, Δ*tatE*) (37), DADE-P (as DADE, *pcn*B1 *zad*-981::Tn*10*d (Km^r^) (38) and MΔBC as MC4100, Δ*tatABC*) (21) were used throughout this study. Strain XL1blue (Agilent) was used for all cloning steps. All plasmids used and constructed in this work are listed in Table 1.

**Table 1.**
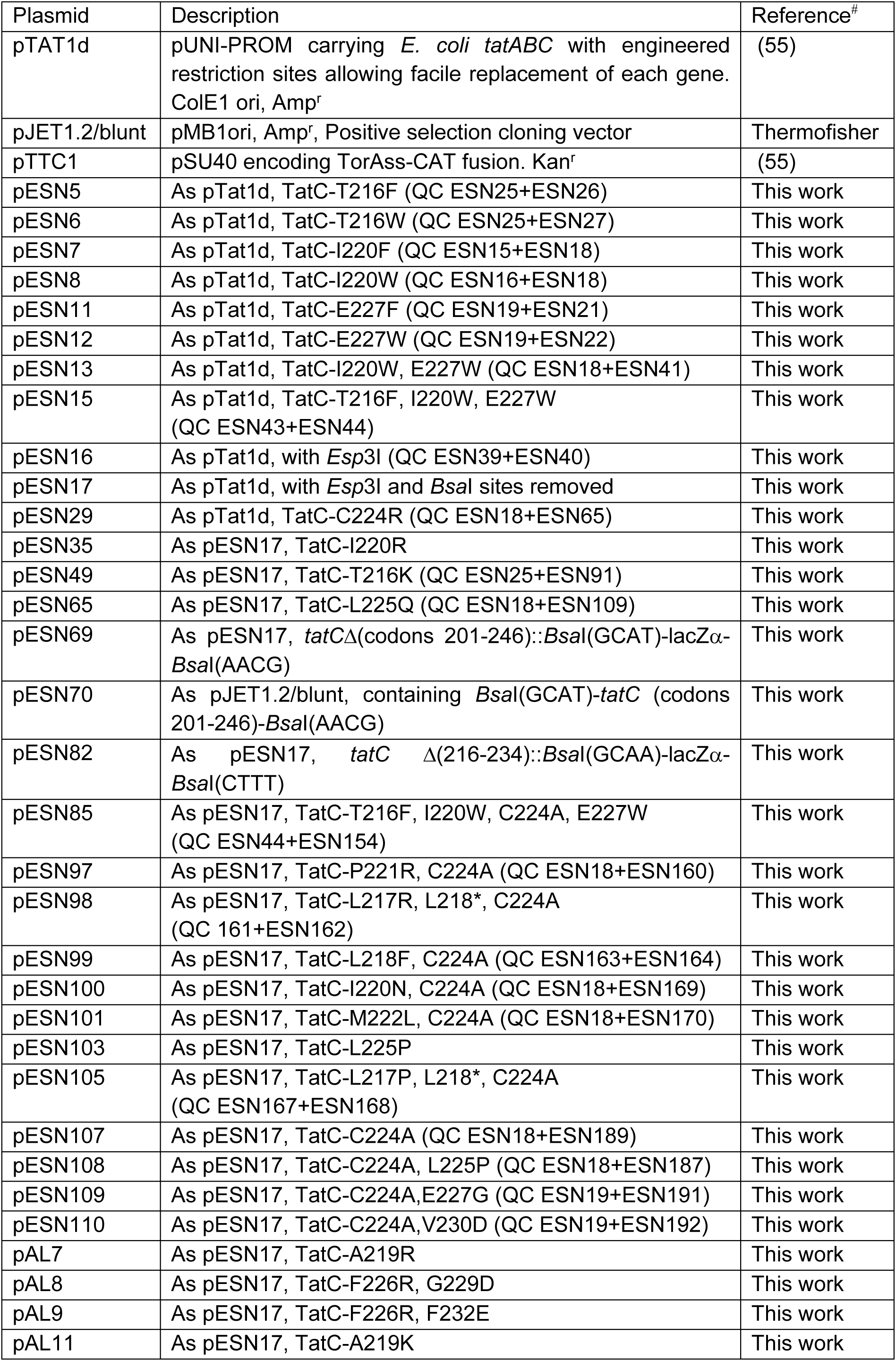
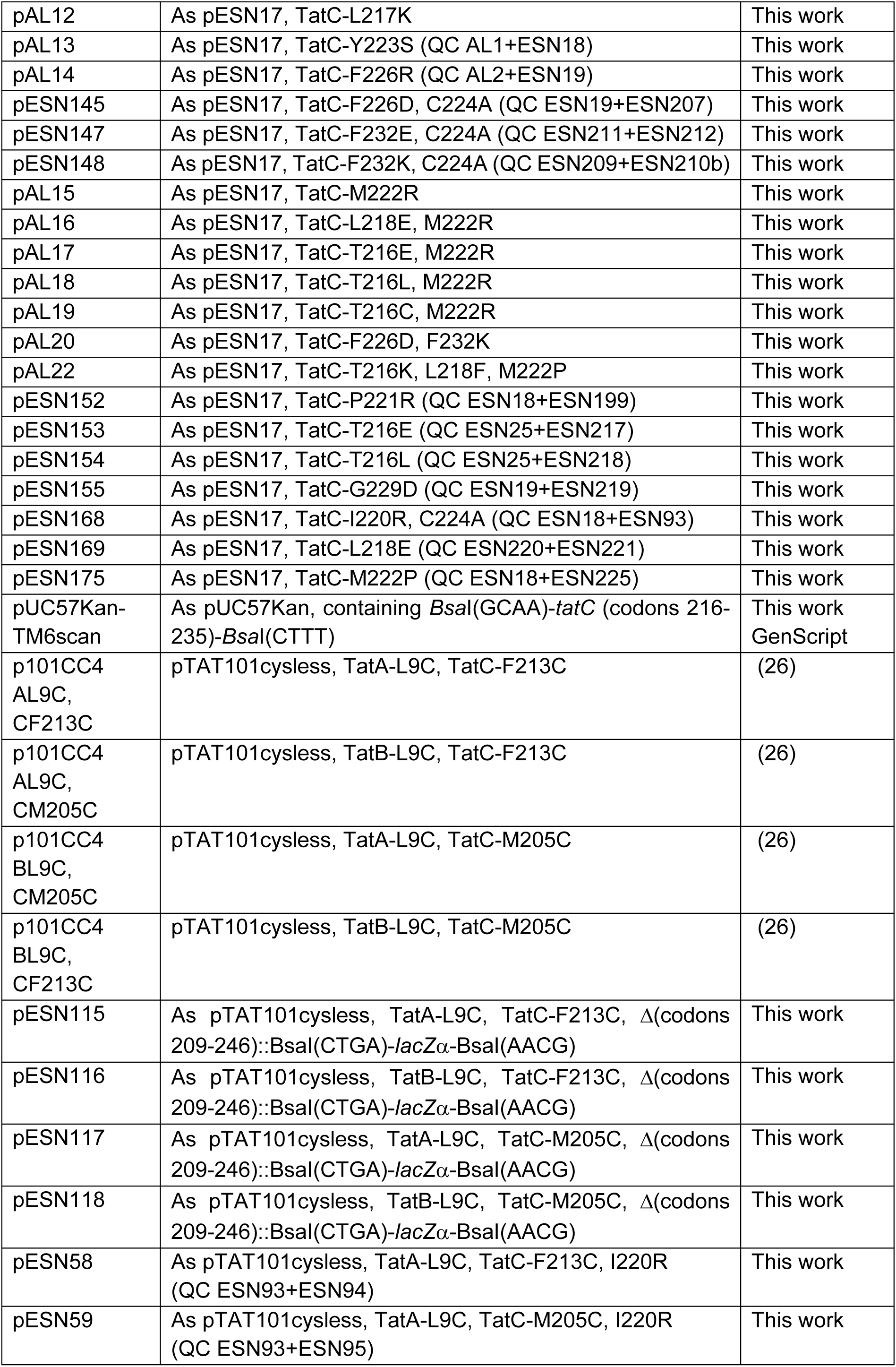
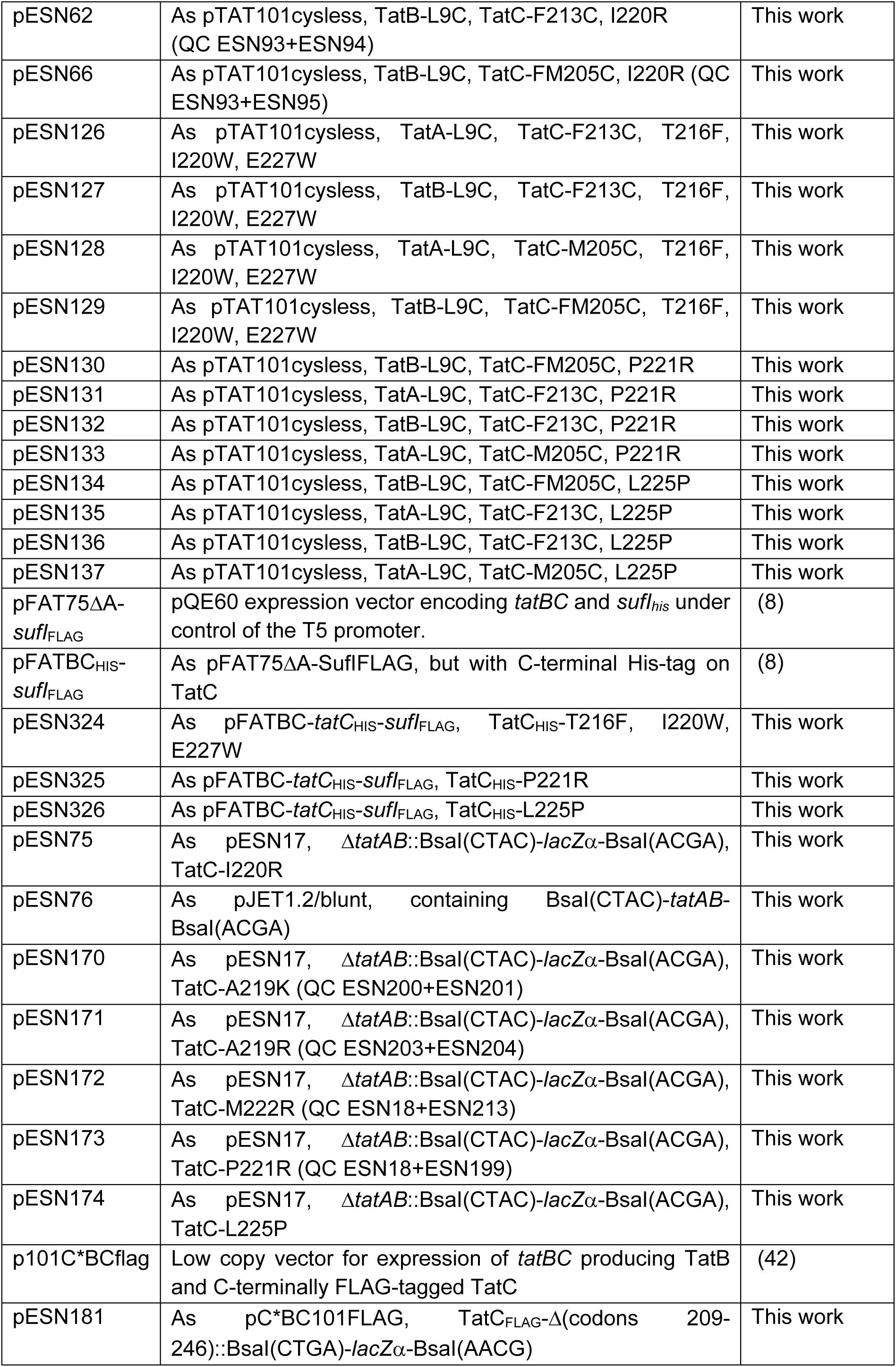
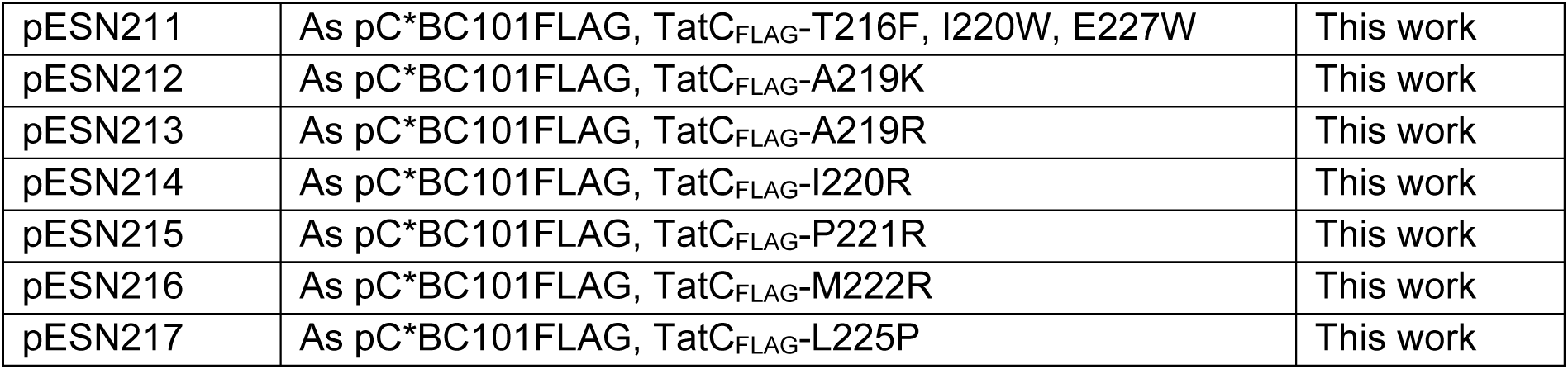
Plasmids used and constructed in this study. For plasmids where substitutions were introduced by QuickChange, these are indicated by parenthesis that includes the names of specific primer pairs used to make them (listed Table S1), preceded by “QC”. Construction details for all other plasmids, which were made by other techniques and/or isolated from library screening, are included in the main Text. *Indicates silent mutation.

### Site-directed mutagenesis

Side-directed mutagenesis of the *tatC* gene was performed with the modified QuickChange^TM^ method described in Liu and Naismith (39). All oligonucleotides used for mutagenesis and cloning and sequencing are listed in Table S1. The templates for whole-plasmid PCR amplification were either pTAT1d/pESN17 (WT *tatABC* template) or other constructs already bearing mutations in the case of multi-site mutants. Finished PCR reactions were treated with *Dpn*I prior to transformation. All constructs were confirmed by full sequencing of the *tat* genes before use.

### Construction of random mutagenesis libraries

Error-prone PCR (ePCR) was used to construct random mutagenesis libraries covering the region encoding TatC TM6. We first constructed a template plasmid, pESN70 (Table 1), by cloning into pJET1.2/blunt (ThermoFisher) a PCR product amplified with primers ESN51+ESN52 (Table S1), encoding TatC TM6 as well as adjacent parts of the TM5-TM6 periplasmic loop and the cytoplasmic C-terminus (codons 201-246). The primers also introduced *Bsa*I sites for Golden Gate assembly. Randomised PCR products were amplified from pESN70 using oligonucleotides ESN118 and ESN119 (Table S1). Reactions were carried out in in 50 μl and contained approximately 50 ng of template DNA alongside 0.2 mM each of dATP/dGTP, 1 mM each of dCTP/dTTP, 7 mM MgCl_2_, 0.4 mM each primer, 5 U GoTaq G2, 1X GoTaq buffer, and a range of MnCl_2_ concentrations (0, 0.1, 0.2, 0.3, 0.4, 0.5, or 0.6 mM) to modulate the mutation rate. The PCR cycle was: 94 C, 2 min; 20x(94 C, 30 sec; 50 C, 30 sec; 72 C, 30 sec); 72, 1min. PCR products were then treated with *Dpn*I for 1-2 hr and then purified (QiaQuick, Qiagen) before use in library construction.

Golden Gate assembly (40) was used to clone each ePCR product into the recipient vector pESN69 (construction details below). Plasmid pESN69 is a modified version of pTAT1d where the *Bsa*I and *Esp*3I sites have been removed by mutagenesis, and where *tatC* codons 201-246 have been replaced with a *lacZ*α cassette flanked by *Bsa*I sites specifying quadruplets <u>GCAT</u> and <u>AACG</u> (codons 200 and 247 underlined) for seamless ligation. One-pot Golden Gate reactions were assembled in 20 μl as follows: ca 250 ng pESN69, 80-100 ng ePCR, 1X T4 ligase buffer, 20 U *Bsa*I-HFv2 (NEB) and 20 U T4 ligase (NEB), and run in a thermal cycler with an initial 1 hr 37 C step followed by: 50x(37 C, 2 min; 16 C, 5 min); 50 C, 5 min; 80 C, 10 min.

Electrocompetent XL1blue cells (Agilent) were electroporated with 1 μl assembly mix and plated onto LB agar containing ampicillin along with 0.25mM isopropyl β-D-1-thiogalactopyranoside (IPTG) and 60 μg/ml 5-Bromo-4-Chloro-3-Indolyl β-D-Galactopyranoside (X-Gal). We obtained between 80 000 and 130 000 colonies per transformation of which none were blue, indicating close to 100% efficiency of assembly. Colonies were harvested in LB medium containing ampicillin, inoculated at an initial OD_600_ of 0.1 into fresh LB medium containing ampicillin and cultured for 5-6 hr before harvesting 3-4 X 5 ml aliquots. Plasmid DNA was isolated from each aliquot, and the aliquots were pooled and used as the final library.

To determine the mutation rate in each library, 10-12 random colonies were picked and plasmid DNA isolated and sequenced (using primer ESN33, Table S1), from which we calculated a figure for the percentage error rate. As the codons targeted for mutagenesis covered approximately 140 nucleotides, we undertook functional screening using the library with a 0.6% error rate (constructed from the ePCR amplified with 0.4 mM MnCl_2_) which yields on average one basepair change in the target DNA.

ePCR was also used to construct random libraries of the *tatAB* bicistron to seek intergenic suppressors of all TatC TM6 mutants. The construction of these libraries used the same design and procedures as described for *tatC*. Briefly, the *tatAB* template plasmid pESN76 was built by cloning a PCR product (ESN74+ESN75, Table S1) into pJET1.2/blunt. This plasmid served as template for ePCR reactions with MnCl_2_ concentrations of 0.1, 0.2, 0.3, and 0.4 mM, and the resulting PCR products cloned into the Golden Gate vector, pESN75 (Table 1-see construction below) which was then moved into XL1blue for sequencing. We undertook functional selection with the library produced with 0.2 mM MnCl_2_, which returned an error rate of ca 0.6 % equivalent to an average of 5 basepair changes in the target DNA (primers ESN45 and ESN140 were used for sequencing). Suppressor libraries for all other inactive *tatC* TM6 mutants were constructed in the isogenic vectors pESN170, pESN171, pESN172, pESN173, pESN174 (see below). All transformations in XL1blue to build the libraries yielded approximately 1 000 000 transformants.

### Scanning mutagenesis library of the *tatC* TM6 coding region

A synthetic scanning mutagenesis library of *tatC* codons 216 – 234 inclusive, comprising nominally 380 single mutants (stop codons were excluded) was synthesised by GenScript in vector pUC57Kan, with the synthetic fragments flanked by *Bsa*I sites. These fragments were moved via Golden Gate assembly (as described above, and using a 1:1 ratio of donor:recipient plasmids) into the recipient construct pESN82 (Table 1; construction details below), and electroporated into electrocompetent XL1blue cells.

### Library screening

The method for isolating inactive *tatC* mutants has been described elsewhere (41). Briefly, mutagenesis libraries were introduced into strain, DADE, pre-transformed with plasmid pTTC1, which encodes chloramphenicol acetyltransferase fused to a twin-arginine signal sequence. Transformants were first selected on LB containing ampicillin and chloramphenicol (the latter at 200 – 400 μg/ml), and then replica-plated onto LB ampicillin and LB ampicillin plus 2%SDS to identify SDS-sensitive clones. Chl^R^ and SDS^S^ clones were patched on LB Amp and preliminarily screened by colony PCR (to quickly eliminate stop codon and frame-shift mutations). Clones harbouring mis-sense mutations were individually re-tested for SDS-sensitivity and, once confirmed, used for plasmid prep and re-transformation of *E. coli* XL1blue to segregate the pTAT1d-derivatives from pTTC1. These constructs were then fully sequenced, and again subjected to phenotypic testing after re-introduction into strain DADE.

Screening of ePCR *tatAB* libraries was performed as above except that libraries were moved into DADE cells and selected directly on 2% SDS LB plates. While the synthetic scanning mutagenesis library of TatC TM6 was designed to harbour only single codon substitutions, it also contained a small proportion of multiple mutants and codon deletions, due to synthesis errors. Inactive *tatC* mutants harbouring multiple substitutions mapping to TM6 (codons 216 - 234) from either library were subsequently deconvoluted by engineering each mutation singularly into pESN17 by QuickChange (see above), followed by individual re-testing of these single mutants to determine whether the individual substitution resulted in TatC inactivation.

### Construction of recipient vectors for Golden Gate assembly

To construct Golden Gate recipient vectors for phenotypic characterisation in DADE, pTAT1d was first modified by mutagenesis to remove the native *Bsa*I and *Esp*3I sites. Initially the *Esp*3I site was removed using Quickchange (ESN39+ESN40) to generate plasmid pESN16. Next, two PCR products were amplified using mutagenic oligonucleotides (Table S1) to suppress the two native *Bsa*I sites remaining in pESN16: oligonucleotides ESN47+ESN48 to remove the *Bsa*I site in *bla* and oligonucleotides ESN49+ESN50 to remove the *Bsa*I site just upstream of the *tat* promoter, respectively. Following digestion with *Dpn*I, these PCR products were Golden Gate-assembled using the engineered *Bsa*I sites present at their ends (Table S1), to generate plasmid pESN17.

The Golden Gate recipient vectors, pESN69 and pESN82, were derived from pESN17 in an equivalent manner using the NEBuilder HiFi DNA assembly kit (NEB). Plasmid pESN69 derived from the assembly of PCR products: ESN114+ESN115 from pESN17 (entire plasmid minus *tatC* codons 201-246) and ESN112+ESN113 on gDNA of *E. coli* MG1655 (*lacZ*α). pESN82 derived from PCR products: ESN152+ESN153 from pESN17 (entire plasmid minus *tatC* codons 216-234) and ESN128+ESN129 from pESN69 (*lacZ*α). Both assemblies were moved into XL1blue and screened on X-Gal-containing media for blue colour. The entire modified *tat* operon in both constructs was fully sequenced and correct cutting by *Bsa*I was confirmed by restriction digestion before further use.

The Golden Gate recipient vectors for the screening of *tatAB* ePCR libraries were made in two steps. First, pESN75 (C-I220R) and pESN174 (C-L225P) (Table 1) were constructed by NEBuilder HiFi assembly similar to that described for pESN69 above, i.e. by combining two PCR products, one for the backbone minus *tatAB* (ESN125+ESN126 on either pESN35 or pESN103, respectively) and one for *Bsa*I-flanked *lacZ*α (ESN123+ESN124 using gDNA of MG1655 as template). Vectors pESN170 - pESN173, harbouring all other inactivating TM6 mutations (Table 1), were then derived from pESN75 by QuickChange (Table S1). All constructs had their modified *tat* operon fully sequenced before use.

To simplify the making of plasmids for *in vivo* disulfide cross-linking (see below), we also constructed Golden Gate recipient vectors based on the previously described set of low-copy pTAT101cysless plasmids carrying unique pairwise combinations of Cys substitutions in either TatC TM6 (F213C) or TM5 (M205C) and either TatA (L9C) or TatB (L9C) (26). Golden Gate recipient constructs, pESN115, pESN116, pESN117, and pESN118 (Table 1), were constructed by NEBuilder HiFi assembly from the following PCR products: ESN128+ESN129 on pESN69 (*lacZ*α) and either ESN139+ESN174 (F213C) or ESN139+ESN175 (M205C) on the appropriate pTAT101cysless templates (entire plasmid minus codons 209-246). The final constructs were screened and confirmed as described above for pESN69 and pESN82.

Finally, we also constructed a Golden Gate recipient vector, pESN181 (Table 1), to facilitate designing plasmids for close to native expression in MΔBC. pESN181 was constructed as described above for pESN115/116 using as the plasmid backbone construct p101C*BCflag (42) which encodes TatB alongside a a C-terminally FLAG-tagged version of TatC. pESN181 was screened and confirmed as described for the other Golden Gate vectors.

### Construction of plasmids for co-purification

Plasmids pESN324, pESN325, and pESN326 are derivatives of pFATBC_HIS_-*sufI*_FLAG_ (42) carrying the TIE (triple T216F, I220W, E227W), P221R, and L225P mutations in *tatC*, respectively. These were obtained using the NEBuilder HiFi DNA assembly kit to join the following PCR products: ESN114+ESN180 from pFATBC_HIS_-*sufI*_FLAG_ (entire plasmid minus *tatC* TM6 coding sequence) and ESN178+ESN182 from either pESN15, pESN152, or pESN103 (Table 1, S1). Inserts were sequenced with forward primer ESN83 to confirm the presence of the correct TM6 mutation as well as that of the histidine tag codons and of the start of *sufI*. As part of the screening process, a plasmid was isolated that carried a single nucleotide deletion at the 5’ junction between the two PCR products, causing a frameshift upstream of the TM6 coding region of *tatC*. This construct, simply named “Δ*tatC*”, was retained as further negative control for the co-purification assays.

### Construction of plasmids for *in vivo* disulfide cross-linking

All constructs for *in vivo* disulfide cross-linking (Table 1) were made in two steps. First, QuickChange was used to introduce the C224A substitution in pTAT1d-based constructs: ESN44+ESN154 on pESN15 to give pESN85 (TIE), ESN18+ESN93 on pESN17 to give pESN168 (I220R), ESN18+ESN160 on pESN17 to give pESN97 (P221R), and ESN18+ESN187 on pESN17 to give pESN108 (L225P) (Table 1, S1). In the second step, these plasmids were used as templates for two sets of PCRs that were Golden Gate-assembled into the final pTAT101cysless-based recipients (see above): ESN132+ESN177 for pESN115 and pESN116 (F213C), and ESN132+ESN176 for pESN117 and pESN118 (M205C). Final constructs (Table 1) were sequenced with primer ESN104 to confirm correct assembly and the presence of the expected TM6 mutation and Cys replacements.

### Construction of plasmids for expression in strain MΔBC

ESN132+ESN176 PCR products amplified from plasmids pESN15, pESN35, pESN103, pESN152, pAL7, pAL11, and pAL15 were Golden Gate-assembled into pESN181 (see above). Final constructs (Table 1) were confirmed as described above for the cross-linking plasmids.

### Preparation of membrane fractions

Membrane fractions of *E. coli* cultures were prepared as described previously (43). Briefly, overnight cultures grown from single colonies of fresh transformants were refreshed in LB supplemented with the appropriate antibiotics (if required) to an initial OD_600_ of 0.05 and grown at 37 °C with shaking until an OD_600_ of 0.5-1. Cells were then harvested from a volume of 25-50 ml (depending on the plasmid copy number), resuspended in 1 ml of the buffer required for each particular application (see below) and supplemented with protease inhibitors (Roche), and then disrupted by sonication. After clarification, membrane fractions were separated from the cell lysate by ultracentrifugation and finally resuspended in 30-60 μl Buffer 2 (50 mM Tris-HCl, pH7.5, 5 mM MgCl_2_, 10% glycerol) unless indicated otherwise.

### Blue-native-PAGE

Blue-Native (BN) PAGE was performed as described (42) except that digitonin-mediated solubilisation was performed overnight. Briefly, membrane fractions were prepared as described above, but with phosphate buffered saline used for cell resuspension and sonication, and Buffer A (50 mM NaCl, 50 mM imidazole, 2 mM 6-aminohexanoic acid, 1 mM ethylenediaminetetraacetic acid/EDTA; pH 7.0) used for final resuspension. After solubilisation of the membrane fractions with 2 % digitonin, the solubilised material was recovered by ultracentrifugation and mixed with 5% glycerol and 0.25% Coomassie blue G-250 buffer Approximately 20 μg of total protein was loaded per lane. Gels were subsequently analysed by western blotting using an anti-TatC antibody (44). Blots were developed with ECL (BioRad).

### Co-purification of TatC_HIS_, TatB, and SufI_FLAG_

For co-purification of TatC_HIS_, TatB, and SufI_FLAG_, cultures of DADE-P carrying either pFAT75ΔA-*sufI*_FLAG_ (control plasmid), or pBC_HIS_-*sufI*_FLAG_, derivatives were grown as for membrane fraction preparation, but in the presence of 1 mM IPTG. After sonication, clarified cell lysates were solubilised overnight with 1.5 % digitonin in Buffer 1 (20 mM Tris-HCl, pH7.5, 200 mM NaCl, 50 mM imidazole, 10% glycerol) and the soluble fraction separated by ultracentrifugation. TatC_HIS_ complexes with TatB and SufI_FLAG_ were captured with Ni-NTA magnetic beads (ThermoFisher), which were washed three times with wash buffer (20 mM Tris-HCl, pH 7.5, 200 mM NaCl, 25 mM imidazole, 0.5% digitonin) before elution with 1X SDS PAGE loading buffer (Brand) by mild thermal treatment (53 °C for 10 min). Samples were then run on SDS PAGE gradient gels (BioRad) and analysed by western blotting with anti-TatB (15), anti-His (Invitrogen), or anti-FLAG (Sigma) antibodies. Blots were developed with ECL (BioRad).

### *in vivo* disulfide cross-linking

*In vivo* disulfide cross-linking was performed as described (26). Briefly, strain DADE harbouring pTAT101cysless-based plasmids (Table 1) were prepared for membrane fractions as outlined above. Once at 0.5 OD_600_, each 50 ml culture was split in two equal volumes which were reacted with either copper phenanthroline (CuP) 1.8 mM or 10 mM 1,4-dithiothreitol (DTT) for 1 min before being washed with ResB buffer (20 mM Tris·HCl, 200 mM NaCl, pH7.5) and finally resuspended in 1 ml ResB supplemented with 12 mM EDTA and 8 mM *N*-ethylmaleimide (NEM) to stop the reaction. Samples were then subjected to membrane fractionation as outlined above. 10 μl/sample were used for SDS PAGE and Western Blotting.

### TatC expression levels in strain MΔBC

Membrane fractions were prepared as described above from strain MΔBC carrying pC*BC101FLAG derivatives (Table 1) using ResB (see above) as wash buffer. 10 μl/sample were used for SDS PAGE and Western Blotting. Monoclonal anti-FLAG antibody M2 (Sigma) was used for detection of TatC_FLAG_.

### Molecular modelling and simulations

Molecular modelling of the TatABC multimer was carried out as described previously (15). Multimers were built using TatA–TatC/TatB–TatC disulfide cross-links as unambiguous constraints for docking using HADDOCK (45). For all models, TatA was modelled from residues G2 to G21, TatB from residues F2 to G21 and TatC from residues T11 to F235. The CHARMM-GUI membrane builder functionality (46) was used to setup the systems for MD simulations. The TatABC complexes (WT and P221R, L225 and TIE mutants) were inserted in a bilayer containing 25% 1-palmitoyl, 2-oleoyl phosphatidylglycerol and 75% 1-palmitoyl, 2-oleoyl phosphatidylethanolamine and solvated with TIP3P waters and 0.15 M NaCl. All MD simulations were performed using GROMACS 2021 (47) and CHARMM36m force field (48)with a timestep of 2 fs. The input files for minimization and equilibration provided by CHARMM-GUI were used. The system was first minimized followed by a series of NVT and NPT equilibration steps consisting of gradual removal of the restraints from lipid and protein atoms for a total time of 2 ns. Three repeats of 100 ns of unrestrained atomistic MD simulations, for each configuration of the molecular system were performed. All simulations were executed at 300 K and 1 bar with protein, lipids and solvent separately coupled to an external bath, using the velocity-rescale thermostat (49) and Parrinello-Rahman (50) barostat. All bonds were constrained with the LINCS algorithm (51). The long-range electrostatic interactions were computed with the Particle Mesh Ewald method (52), while a Verlet cut-off method was used to compute the non-bonded interactions. All images were generated using PyMOL (53).

## RESULTS

### Bulky substitutions at the TatA/TatB binding site on TatC TMH6 do not abolish Tat function

Structural analysis of *Aquifex aeolicus* TatC identified that TMH5 and TMH6 contain fewer amino acids than conventional TM helices, and therefore are slightly shorter in length than normal TM helices. This region of TatC proteins comprises a number of highly co-evolving residue pairs, which suggests tight structural interplay between both TMHs (9). A primary binding site for TatA/TatB has been characterised along TMH5 of TatC, with residues in the adjacent loop and the top of TMH6 also forming a key part of the site; referred to as the polar cluster site (Fig 2A). The TMH6 binding site is largely formed by residues in TMH6, including F213, I220 and E227 (Fig 2A,B, all numbering is for *E. coli* TatC). Initially we took a targeted approach in an attempt to disrupt the TatA/TatB binding site on TatC TMH6. We selected residues that fall on the face of TatC TMH6, which have been shown to interact with TatA by crosslinking studies (26) (Fig 2B). We also mutated each of T216, I220 and E227 singly and in combination to bulky phenylalanine (T216F) or tryptophan (I220W, E227W). We then tested for Tat transport activity by plating cells producing these variants of TatC onto LB containing 2% SDS. Complete block of the Tat pathway results in the failure to export the two cell wall amidases, AmiA and AmiC, to the periplasm and the resultant defect in cell wall remodelling makes cells exquisitely sensitive to killing by detergent (54). However, as seen in Fig 1C, even a TatC variant where T216, I220 and E227 are triply-substituted (‘TIE’ mutation) did not abolish Tat activity and was able to support robust growth on SDS.

**Fig 2.**
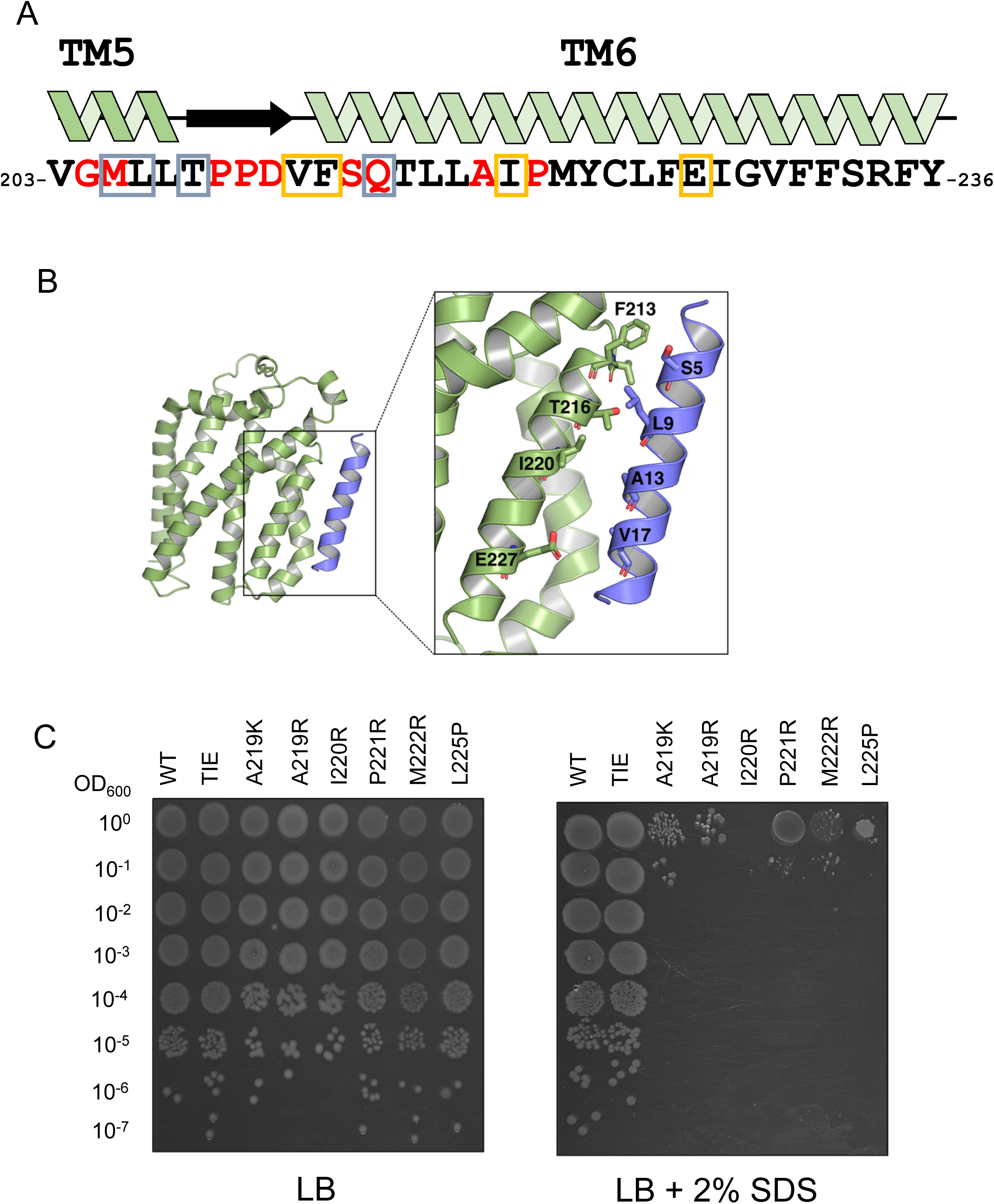
Mutagenesis of TatC TMH6. (a) amino acid sequence of TatC TMH6 and flanking regions. Inactivating substitutions falling in this region of TatC that have been identified previously (15, 41, 57) are indicated by red font. Residues shown in grey box have been shown by cysteine-substitution and crosslinking (M205, L206; (26)) or co-evolution analysis, molecular simulations, and mutagenesis (15) to form the TatA/TatB TMH5 binding site. Residues shown in yellow box are part of the TMH6 binding site (26). (b) Structural model of TatA bound at the TMH6 binding site. (c) Phenotypic characterisation of TatC single-amino acid TM6 variants. Spot tests of strain DADE (Δ*tatABCD*, Δ*tatE*) carrying pTAT1d derivatives producing TatA, TatB and the indicated amino-acid TM6 variant in TatC. Strains were re-grown from overnight cultures in liquid medium to 1 OD600, decimally diluted and spotted (10 ml) on LB medium with or without added 2% SDS, and finally incubated overnight aerobically at 37 °C.

### Random mutagenesis of the TatC TMH6 coding region

Next, we turned to random mutagenesis as an unbiased approach to isolate inactivating amino acid substitutions in TatC TMH6. To date, only two inactivating substitutions, A219E and L225P have been reported (41). To identify further inactivating substitutions, we initially used error-prone PCR to construct a random library of 90,000 clones containing mutations covering the coding region for TatC TMH6 and flanking regions, as described in Methods. To isolate amino acid exchanges that compromise the activity of TatC, we used a 2-step approach that we have previously successfully employed to identify inactivating substitutions in Tat components (41). The TorA-CAT reporter protein comprises the twin-arginine signal sequence of the Tat substrate protein TorA, fused to chloramphenicol acetyltransferase (55). This fusion protein is exported to the periplasm of a wild-type strain, rendering cells sensitive to growth inhibition by chloramphenicol, which can only be inactivated by CAT in the cytoplasmic compartment. Substitutions that reduce or abolish Tat transport allow cytoplasmic CAT to accumulate, conferring resistance to chloramphenicol. As this initial screen does not differentiate substitutions that reduce Tat activity from those that fully abolish it, all chloramphenicol-resistant colonies from the first round of screening were subsequently plated onto LB containing 2% SDS to identify substitutions that completely abolish TatC activity.

Following extensive screening of the library we found 101 mutant clones that inactivated the function of TatC. Of those, 53 contained a premature stop codon and 10 had frameshifts (Table S2). The amino acid substitutions that were present in the remaining 38 clones are listed in Table 2. A number of substitutions were isolated that fell in the C-terminal region of TMH5 and the TMH5-TMH6 loop region, including M205K/R and Q215R, that have been isolated previously (41). We also isolated seven clones harbouring the TatC L225P substitution, including three where it was the sole mutation present. For the clones with multiple substitutions, we followed up by making the single mutations L217P, L217R, I220N, P221R, M222L, Y223S, C224R, L225Q, E227G and V230D. From this deconvolution, P221R was the only further single substitution that inactivated TatC.

**Table 2.** Inactivating amino acid substitutions in TatC isolated from an error-prone PCR library covering TM6. ‘–’ indicates that the clones conferred full sensitivity to SDS (as determined by the inability of a 10μl of culture of an OD_600_ of 1 to grow on solid medium containing 2% SDS). ‘+/−‘ indicates that clones conferred partial sensitivity to SDS (as determined by an inability to support growth beyond a 10^−4^ dilution of an OD_600_ = 1 culture). Substitutions shown in bold have been isolated previously and shown to inactivate the function of Tate (41). Those in underline indicate that changes at these amino acid positions are known to inactivate Tate when substituted to something other than that identified here.

| TatC substitution | Growth on SDS |
| --- | --- |
| **M205K**, <u>L225Q</u> | +/- |
| <u>M205T</u>, L207P, <u>D211V</u> | +/- |
| <u>Q215K</u>, I220V | +/- |
| <u>G204W</u>, L217P | +/- |
| L217R, F232S | +/- |
| I220N, M222L, E227G, V230D | +/- |
| L217P, M222L, E227V | +/- |
| <u>D211V</u>, C224R, K239E | +/- |
| P210T, Y223S, C224R | +/- |
| V203I, L207Q, T208A, F213Y, C224R | +/- |
| <u>M205V</u>, L206S, P221R | - |
| V203A, <u>G204R</u> | - |
| <u>G204R</u>, I228V | - |
| V203D, <u>G204R</u>, V212A, L218K, F226L | - |
| **M205R** | - |
| **M205R**, G229S | - |
| **Q215R** (isolated 5 times independently) | - |
| **Q215R**, Y236H | - |
| V212I, **Q215R** | - |
| <u>M205L</u>, **Q215R** | - |
| **Q215R**, K239N | - |
| V212A, **Q215R**, Y223H, V230D | - |
| <u>D211E</u>, **Q215R**, T216K, L217Q | - |
| <u>D211G</u>, **Q215R** | - |
| <u>M205I</u>, **Q215R** | - |
| **Q215R**, F235S | - |
| **Q215R**, C224G, E227D, F235L | - |
| **L225P** (isolated 3 times independently) | - |
| M222V, **L225P**, E227G | - |
| **L225P**, E227G, K239I, E244G | - |
| M222T, **L225P**, F235S, G240W | - |
| F213Y, **L225P** | - |

### A scanning mutagenesis library permits identification of further inactivating substitutions in TatC TMH6

While error prone PCR is a useful approach to generate point mutations, not all substitutions are likely to be covered because some amino acid exchanges require two base changes at a position, we had a commercial library synthesised that encoded every single amino acid change at TatC residues 216 – 234, excluding stop codons. We opted to start the mutagenesis from residue 216 since some amino acids prior to this position also play key roles in the TMH5 binding site. Screening this library identified further substitutions that inactivate TatC (Table 3), including positively charged substitutions at A219 and arginine substitutions at I220 and M222. Fifteen frame shifts and a small number of multiple substitutions were also isolated that represent synthesis errors and are present at very low levels in the library but are amplified in our screen because they inactivate TatC. From the small number of multiple substitutions, we constructed individual TatC T216L, T216K, T216E, L218E, L218F, M222P, F226R, F226D, G229D, F232K and F232E substitutions, all of which retained activity. A T216C substitution has been constructed previously and also shown to be functional (26).

**Table 3.** Inactivating amino acid substitutions in TatC isolated from a scanning mutagenesis library of TM6. ‘–’ indicates that the clones conferred full sensitivity to SDS (as determined by the inability of a 10μl of culture of an OD_600_ of 1 to grow on solid medium containing 2% SDS). ‘+/−‘ indicates that clones conferred partial sensitivity to SDS (as determined by an inability to support growth beyond a 10^−4^ dilution of an OD_600_ of 1 culture) (typically, sensitivity showed between dilutions of −2 and −4).

| TatC substitution | Growth on SDS |
| --- | --- |
| L217K (isolated 2 times) | +/- |
| I220R (isolated 6 times) | - |
| A219R (isolated 6 times) | - |
| A219K (isolated twice) | - |
| M222R | - |
| T216E, M222R (isolated twice) | - |
| T216L, M222R | - |
| T216C, M222R | - |
| L218E, M222R | - |
| F226D, F232K | - |
| F226R, G229D | - |
| F226R, F232E | - |
| T216K, L218F, M222P | - |

Together our mutant screens identified A219K, A219R, I220R, P221R, M222R and L225P as single substitutions inactivating TatC, and Fig 1C shows the SDS growth phenotype associated with each of these substitutions.

Previously we have used suppression genetics to isolate mutants in *tatB* that restore activity to inactivating substitutions in TatC, providing mechanistic insight into their functions (41, 42). We therefore designed mutant libraries of *tatAB* by error prone PCR and used these in attempts to suppress each of our inactive TatC substitutions, selecting for growth on LB containing SDS. However, despite numerous rounds of screening we were unable to isolate any robust suppressors for any of these substitutions. Moreover, the strong TatB suppressor, TatBF13Y which restores Tat activity to substitutions in either TatC signal peptide binding site, or to inactive Tat signal peptides (42), was unable to restore detectable activity to any of our TatC mutants.

### Positively charged amino acids at residues 219 and 220 destabilise TatC

We next biochemically characterised the Tat system harbouring the inactivating substitutions. We first asked whether any of the mutations in TMH6 destabilised TatC. Fig 3 indicates that the A219K, A219R and I220R variants of FLAG-tagged TatC are indeed unstable and cannot be detected in membrane fractions, likely explaining their lack of activity. When we expressed these substitutions from a higher copy number vector, we again observed that these alleles are destabilised relative to wild type TatC (Fig 4B), although we were able to faintly detect some TatC from these constructs.

**Fig 3.**
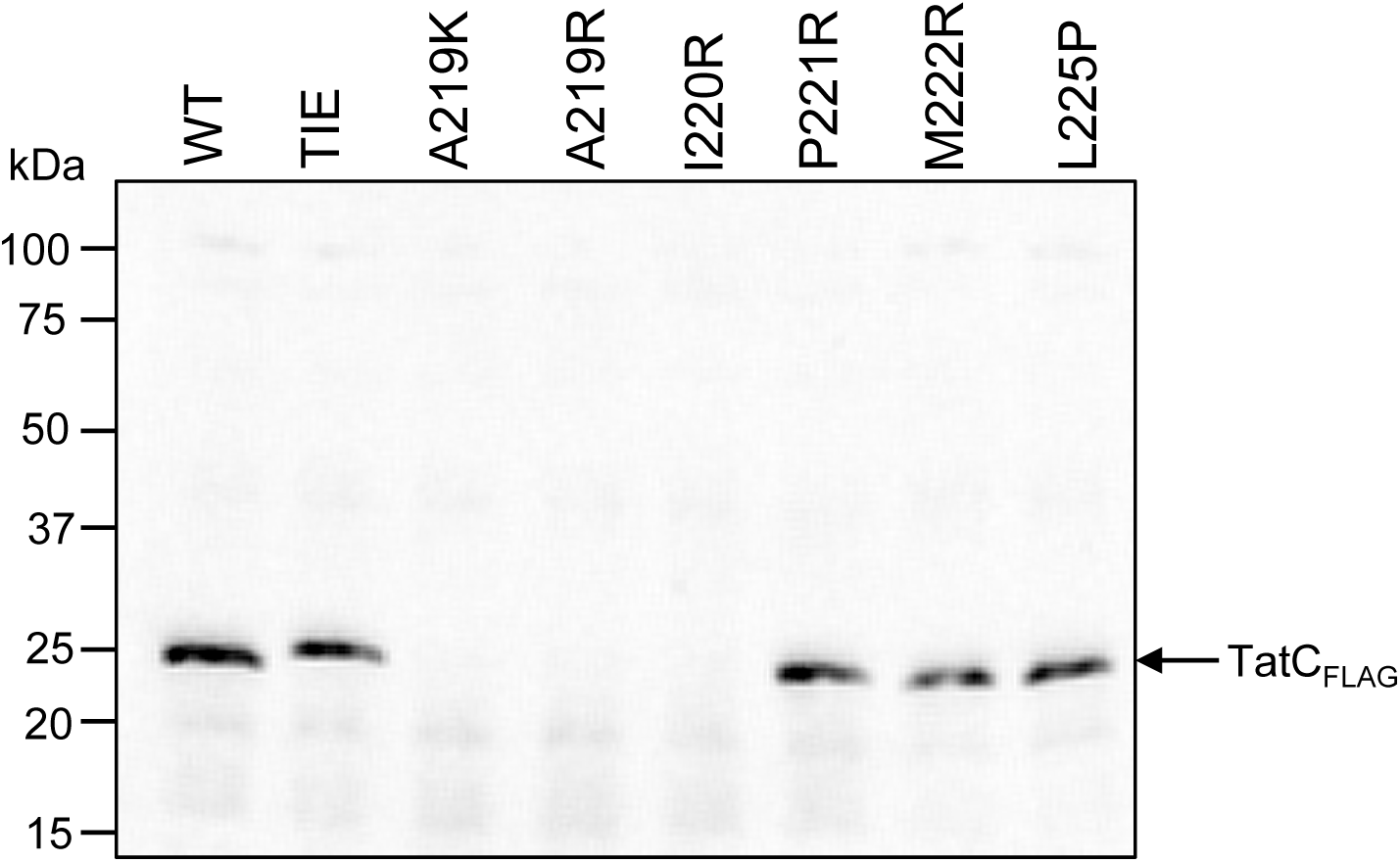
Membrane accumulation levels of TatC single-amino acid TM6 variants expressed from a low-copy vector. Western blots of membrane fractions from MΔBC (as MC4100, Δ*tatBC*) cultures harbouring derivatives of the low-copy construct pC*BC101FLAG expressing TatB and the indicated amino-acid TM6 variant in C-terminally FLAG-tagged TatC. Crude membrane fractions were prepared from exponentially growing cultures as described in Methods, blotted and probed with an anti-FLAG monoclonal antibody. An equal amount of total protein was loaded in each lane.

**Fig 4.**
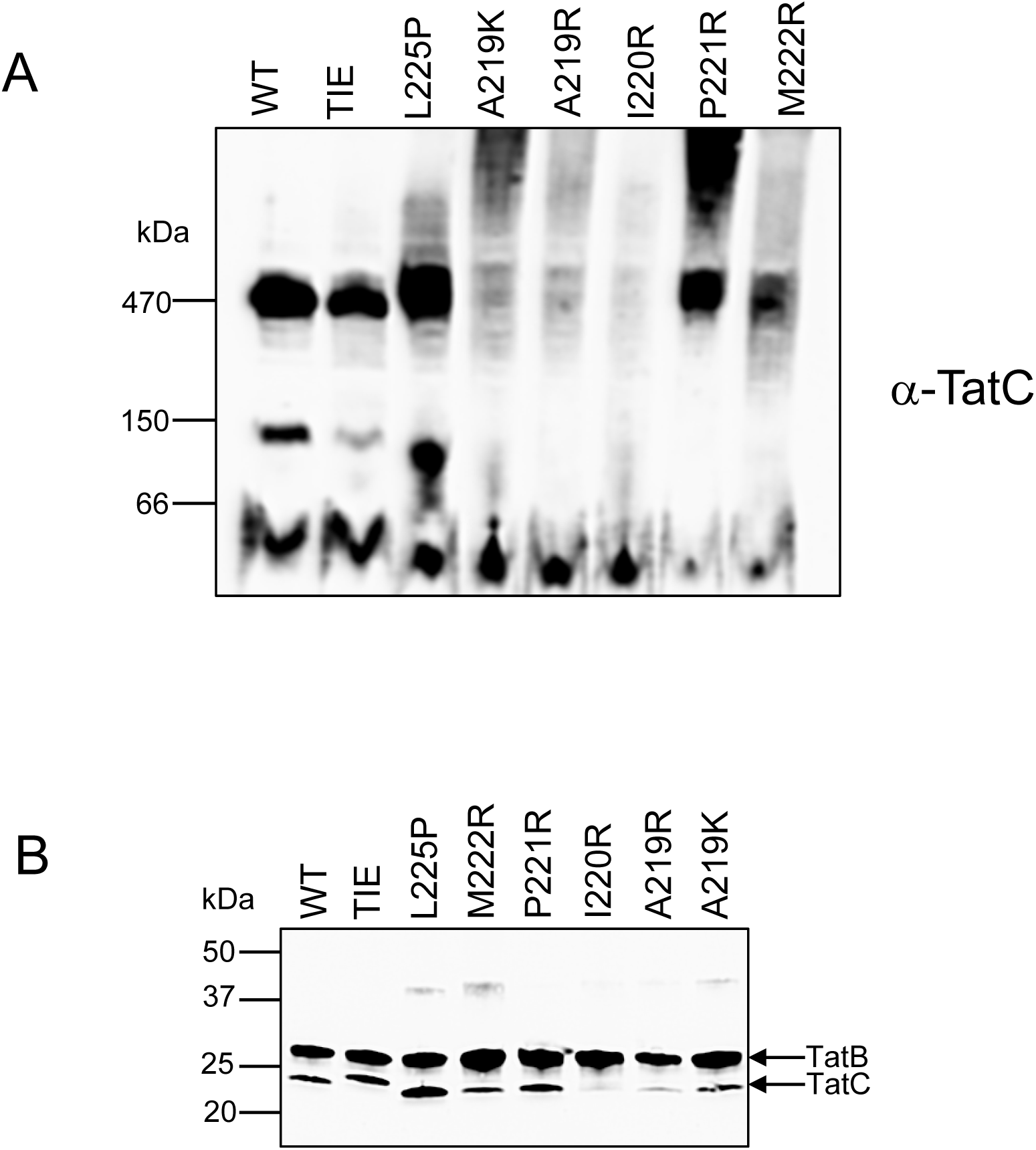
BN-PAGE analysis of Tat complexes containing TatC TMH6 variants. (a) Crude membranes from strain DADE (Δ*tatABCD*, Δ*tatE*) producing the indicated TatC variants alongside wild-type TatA and TatB from plasmid pTAT1d were solubilised by addition of 2% (wt/vol) digitonin and analysed by BN-PAGE (4–16% Bis-Tris NativePAGE gels; 20 μg solubilised membrane/lane) followed by Western blot with an anti-TatC antibody as indicated (b). Crude membranes were also analysed by SDS-PAGE and blotted with a mix of anti-TatB and anti-TatC antibodies as a control to assess protein levels.

The Tat receptor complex has been extensively characterised by blue-native gel electrophoresis, and shown to migrate just above 440kDa on 4-16% gradient gels (e.g. (56)). We therefore tested whether any of the TMH6 substitutions affected migration of the complex. Fig 4A shows that a TatC-reactive complex with a size of around 470kDa was detected when TatC was wild type, and also when it harboured the TIE triple tryptophan substitution (which does not abolish Tat function) or any of the P221R, M222R or L225P inactivating substitutions. As expected, given the lack of stability of TatC A219K, A219R and I220R, no TatC-containing complexes could be detected following BN-PAGE. We conclude that the P221R, M222R and L225P substitutions of TatC do not affect assembly of the Tat receptor.

### TatBC complexes containing the P221R and L225P TatC substitutions can still interact with a Tat substrate

A key function of the Tat receptor complex is the interaction with substrate proteins, mediated through binding with their twin-arginine signal peptides. We therefore sought to determine whether the P221R and L225P TatC substitutions were inactive because they prevented interaction with the Tat substrate, SufI. We co-produced TatB and C-terminally his-tagged TatC alongside SufI-FLAG, isolated membrane fractions, solubilised with detergent and pulled out TatC-his complexes. Fig 5 shows that FLAG-tagged SufI specifically co-purified with TatB and his-tagged TatC, even when the P221R or L225P substitutions were present, and

**Fig 5.**
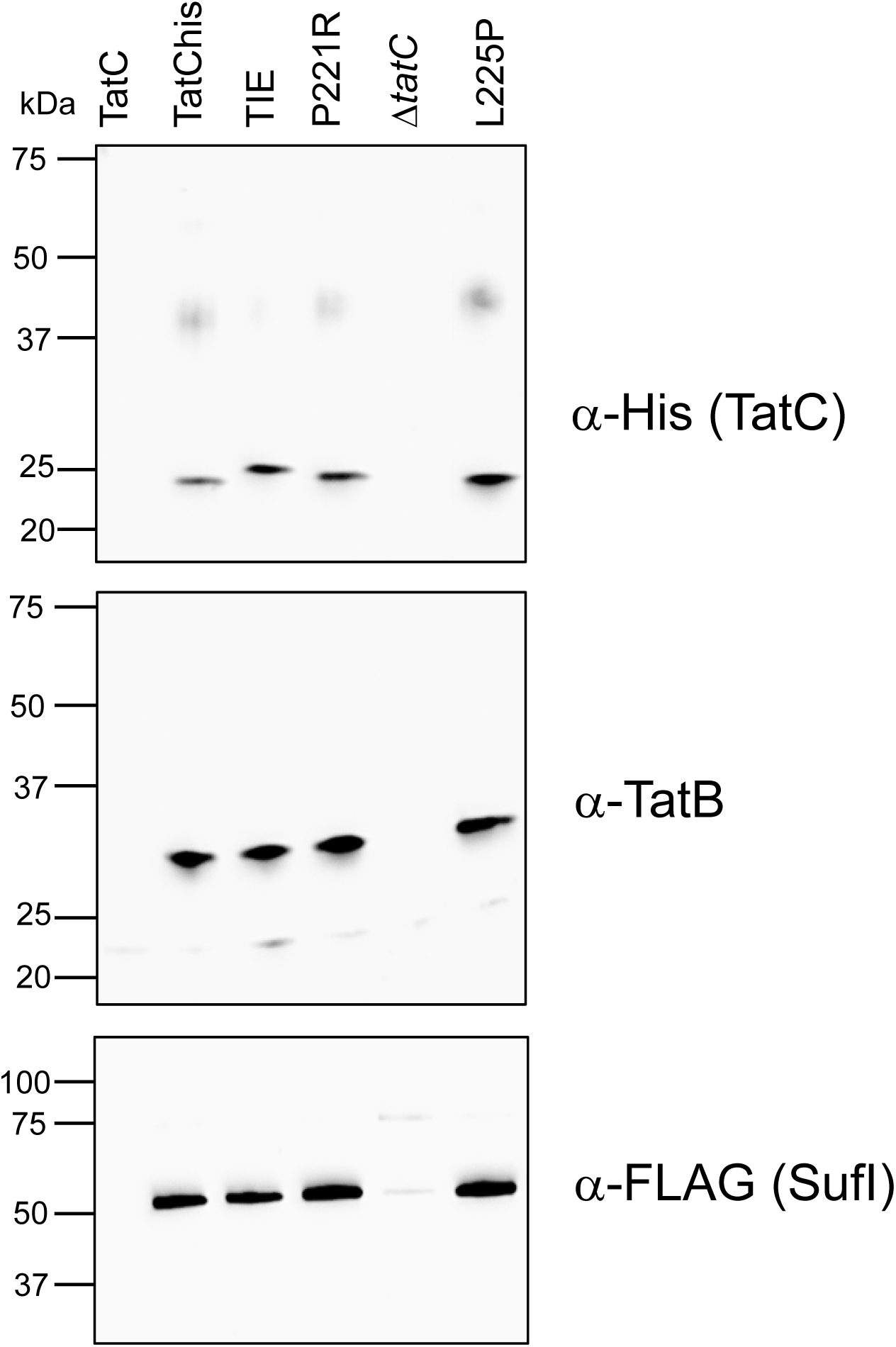
Co-purification of TatB and variant TatC with the Tat substrate SufI. Strain DADE-P (Δ*tatABCD*, Δ*tatE*, *pcnB1*) producing the indicated TatC_HIS_ variants alongside wild-type TatB and the Tat substrate SufI_FLAG_ from plasmid pFATBC_HIS_-*sufI*_FLAG_ were inoculated from overnight cultures at starting of OD600 of 0.05 and grown for 3 hours in the presence of 1 mM IPTG after which membrane fractions were produced as described in Methods. Membranes from were isolated, solubilised by addition of 2% (wt/vol) digitonin and incubated with Ni-NTA-magnetic beads to separate TatC_HIS_-containing complexes. Affinity-bound complexes were eluted by the thermal treatment of the beads and analysed on SDS-PAGE with anti-His (for TatC), anti-TatB, and anti-FLAG antibodies. “*tatC*”: DADE-P harbouring pFATΔA-*sufI*FLAG expressing WT TatC without a His-tag (8); “D*tatC*”: DADE-P carrying a pFATBC_HIS_-*sufI*_FLAG_ derivatives with a PCR-induced frameshift within the TM6 coding region of *tatC*_HIS_, used here as further negative control.

### Disulfide crosslinking reveals that inactivating TMH6 substitutions do not prevent binding of TatA or TatB to the TMH5 or TMH6 sites

Finally, we tested whether any of the inactivating I220R, P221R or L225P substitutions, or the functional ‘TIE’ triple tryptophan substitution of TatC blocked the ability of TatA or TatB to occupy either of the two binding sites. The occupancy of these sites can be probed using site-specific crosslinking; the presence of TatA or TatB in the TMH5 site is shown by a disulfide crosslink between an L9C substitution in TatA or TatB and an M205C substitution in TatC (Fig 6A). Conversely, the presence of TatA/TatB in the TMH6 site is detected through a crosslink between the same L9C substitutions and an F213C substitution in TatC (Fig 6B).

**Fig 6.**
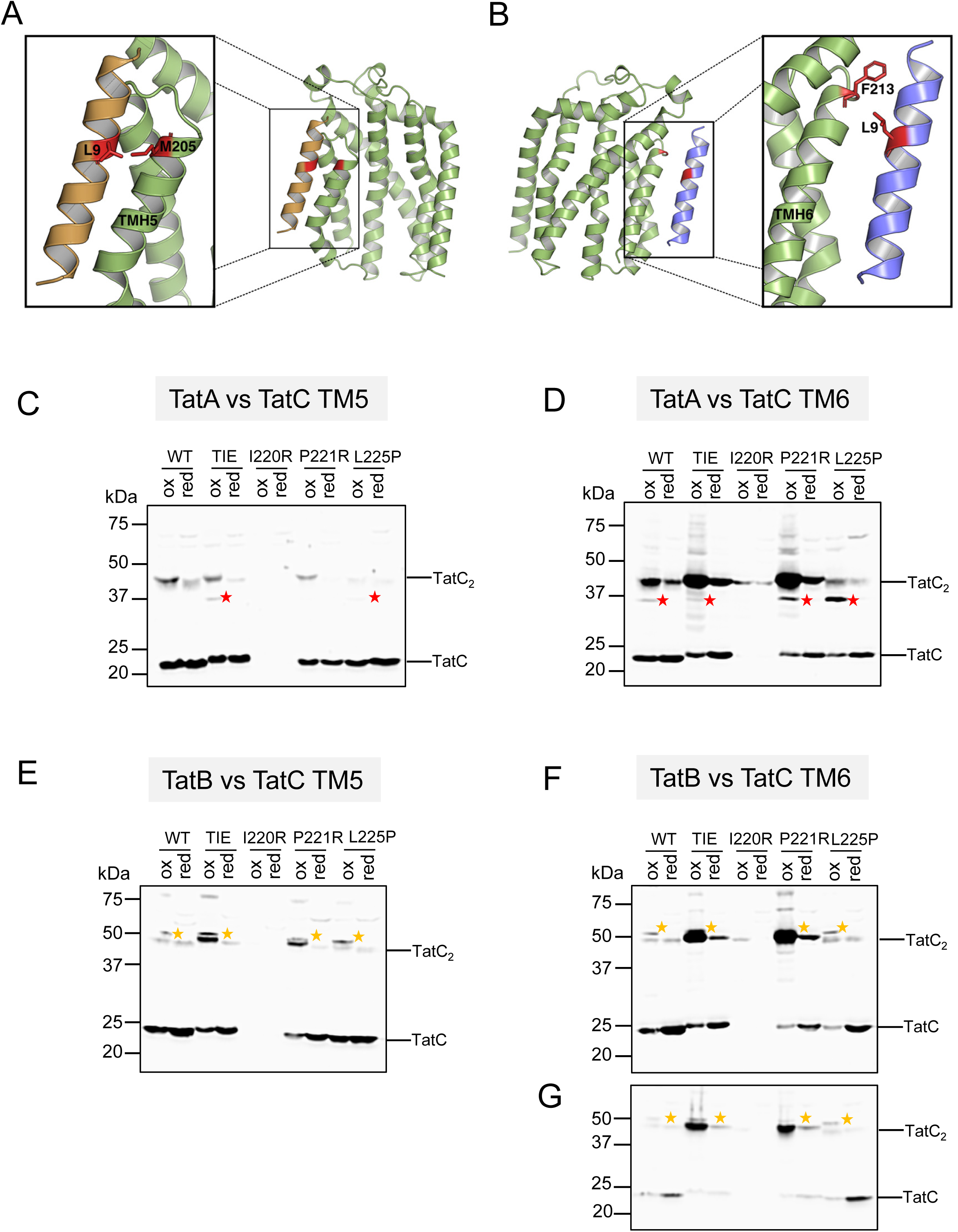
*In vivo* disulfide cross-linking between TatC harbouring TM6 substitutions and TatA or TatB. (A and B). Diagnostic crosslinks between TatA/TatB and TatC can be used to probe occupancy of (A) the TMH5 site (L9C substituted TatA or TatB and M205C TatC) and (B) the TMH6 site (L9C substituted TatA or TatB and F213C TatC). (C-G) Membranes from strain DADE (Δ*tatABCD*, Δ*tatE*) producing the indicated TatC variants <u>and</u> either the M205C or F213C substitutions alongside L9C variants of either TatA or TatB from the low-copy plasmid pTAT101cysless were analysed by anti-TatC western blotting after exposure of whole cells to either 1.8 mM CuP (oxidising, “ox”) or 10 mM DTT (reducing, “red”). Crosslinks are shown between (C) TatA[L9C] and TatC[M205C]; (D) TatA[L9C] and TatC[M213C]; (E) TatB[L9C] and TatC[M205C]; (F) TatB[L9C] and TatC[F213C]; (G) Is reload of the membranes from panel (F) with reduced exposure to prevent signal saturation. In each case red asterisks mark the positions of TatA-TatC crosslinks and yellow asterisks the TatB-TatC crosslinks.

Comparing Fig 6 panels C and D indicates that when TatC is otherwise wildtype, TatA is largely detected in the TMH6 binding site, as reported previously (26). As expected, the I220R substitution resulted in TatC instability and barely any protein could be detected. However, for both the P221R and L225P substitutions, TatA is present at the TMH6 site, indicating that this binding site has not been grossly disrupted by the substitutions. Conversely, very little TatA is detected in the TMH6 site when the triple tryptophan substitutions (‘TIE’) are present. However, this binding site cannot be completely disrupted by these mutations because we could detect some TatB apparently bound there (Fig 6F, G). Likewise, neither the P221R or L225P substitutions completely abolished the ability of TatB to interact with the TMH5 or TMH6 sites. Finally, we note that while there is some TatA at the TMH5 site for the L225P substitution, we could not detect TatA in this site when the TatC P221R substitution was present.

### Molecular Dynamics (MD) Simulations of TatABC complexes

To structurally test the impact of the mutations on modelled complexes of *E. coli* TatABC, we ran three repeats of MD simulations of wild-type, P221R, L225P and TIE variants of TatC, with TatA in the TMH6 binding site and TatB in the TMH5 binding site. For the wild-type simulations both single-pass TMHs remained bound to their respective sites over the course of the simulations, with some deviation observed for TatA bound to TMH6. However, at the TMH6 site, all mutant simulations showed a greater separation between TatA and TatC, with the bulky-substituted TIE variant showing the greatest separation (Fig 7A), in agreement with the very weak TatA L9C-TatC F213C crosslinking we observed when these mutations were present (Fig 6D). Snapshots of the MD simulations, shown in Fig 7C, indicate that for all three of the TatC variants there is TatA disassociation from the TMH6 site, whereas in contrast, TatB remains bound to the TMH5 site for both wild-type and mutant simulations.

**Fig 7.**
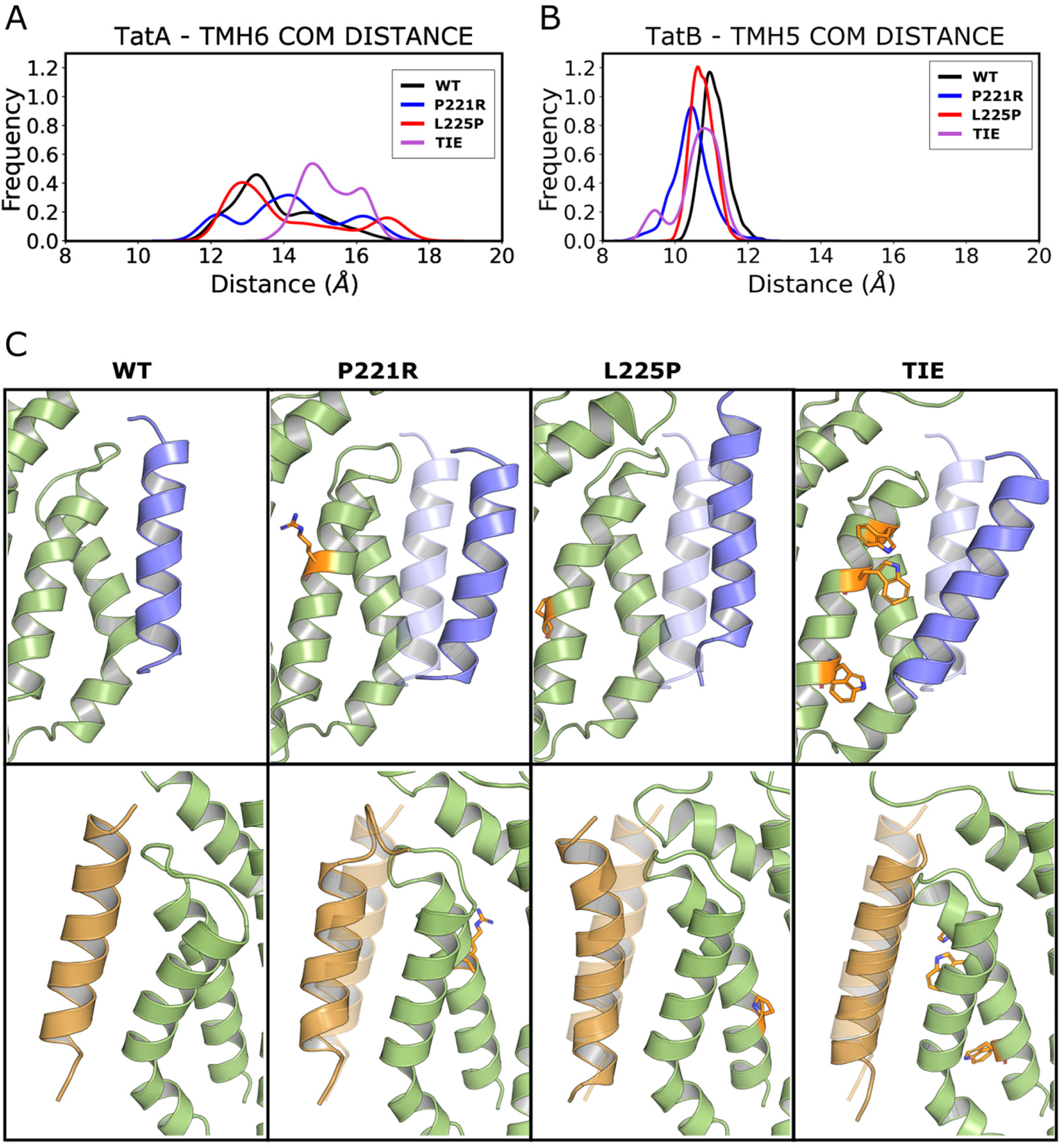
Molecular simulations of the interactions of TatA with TatC TMH6 for the WT and P221R, L225P and TIE mutants. (a) Distances between TatA and TatC TMH6. (b) Distances between TatB and TatC TMH5. The center of mass of the transmembrane helices was used to calculate the distances. (c) Snapshots of the MD simulations showing the displacement of TatA (blue) and TatB (orange) from the Tate (green) TMH6 and TMH5 binding sites, respectively. The three Tate mutants are shown in comparison with the *\NT*.

## DISCUSSION

In this study we have probed the role of TatC TMH6 for function of the Tat pathway. Prior work has shown that the N-terminal end of TMH6 contributes to a key binding site for TatA/TatB, and that this site (here termed the TMH5 site) is mechanistically essential for Tat activity (15, 26, 41). The opposing face of the helix forms a separate interaction interface for TatA/TatB, the TMH6 site, the relevance of which is unclear (26). In an effort to determine whether this binding site is critical for Tat transport, we undertook extensive mutagenesis of TMH6 to identify substitutions that blocked activity. From this we identified six single substitutions that rendered TatC inactive, of which only three produced stable TatC protein. These three TatC variants P221R, M222R and L225P did not affect assembly of the Tat receptor complex, and we confirmed that two of those, P221R and L225P, also did not affect the ability of the receptor to bind a Tat substrate. Using site-specific cross-linking, we found that the P221R and L225P substitutions did not appear to affect the ability of TatA or TatB to interact with the TMH5 or TMH6 binding sites, although MD simulations suggested that TatA binding at the TMH6 should be partially destabilised by either of these substitutions. At present, the explanation for the inactivity of these variant TatC proteins is unclear. It may be that they are unable to undergo conformational changes associated with substrate transport, however we currently lack biochemical tools to address this further.

In a deliberate attempt to disrupt the TMH6 binding site, we also introduced three bulky amino acid sidechains at positions that contact bound TatA (26). MD simulations indicated that these three substitutions would be expected to disrupt interaction with TatA, and indeed site-specific crosslinking revealed that very little TatA could be detected in this site. However, unlike the P221R, M222R and L225P substitutions, this bulky substituted TatC variant did not noticeably affect Tat transport activity. This finding might suggest that the TMH6 site is dispensable for Tat function. However, it should be noted that TatA is in relatively high abundance in *E. coli* membranes (14, 23), and that the TMH6 binding site sits on the outside of the multimeric receptor complex, in close proximity to the membrane (26). It is therefore possible that despite the reduced affinity of TatA for this mutated binding site, transient interaction is sufficient to support Tat transport.

## ACKNOWLEDGEMENTS

This work was supported by the UKRI Medical Research Council through grant MR/S009213/1 to TP and PJS. Research in PJS’s lab is also funded by Wellcome (208361/Z/17/Z) and BBSRC (BB/P01948X/1, BB/R002517/1 and BB/S003339/1). This project made use of time on ARCHER2 and JADE2 granted via the UK High-End Computing Consortium for Biomolecular Simulation, HECBioSim (http://hecbiosim.ac.uk), supported by EPSRC (grant no. EP/R029407/1). This project also used Athena and Sulis at HPC Midlands+, which were funded by the EPSRC on grants EP/P020232/1 and EP/T022108/1. We thank the University of Warwick Scientific Computing Research Technology Platform for computational and Dr Felicity Alcock and Prof Ben Berks for helpful discussion.

## Notes

The authors declare no conflict of interests

### Competing Interest Statement

The authors have declared no competing interest.

## REFERENCES

1. Palmer T, Stansfeld PJ. Targeting of proteins to the twin-arginine translocation pathway. Mol Microbiol 2020;113:861–871.

2. Theg SM. Chloroplast transport and import. Photosynth Res 2018;138(3):261–262.

3. Berks BC. The twin-arginine protein translocation pathway. Annu Rev Biochem 2015;84:843–864.

4. Cline K. Mechanistic Aspects of Folded Protein Transport by the Twin Arginine Translocase (Tat). J Biol Chem 2015;290(27):16530–16538.

5. De Geyter J, Smets D, Karamanou S, Economou A. Inner Membrane Translocases and Insertases. Subcell Biochem 2019;92:337–366.

6. Cristobal S, de Gier JW, Nielsen H, von Heijne G. Competition between Sec- and TAT-dependent protein translocation in *Escherichia coli*. Embo J 1999;18(11):2982–2990.

7. Stanley NR, Palmer T, Berks BC. The twin arginine consensus motif of Tat signal peptides is involved in Sec-independent protein targeting in *Escherichia coli*. J Biol Chem 2000;275(16):11591–11596.

8. Huang Q, Palmer T. Signal Peptide Hydrophobicity Modulates Interaction with the Twin-Arginine Translocase. MBio 2017;8(4).

9. Rollauer SE, Tarry MJ, Graham JE, Jaaskelainen M, Jager F, et al. Structure of the TatC core of the twin-arginine protein transport system. Nature. 2012;492(7428):210–214.

10. Sargent F, Bogsch EG, Stanley NR, Wexler M, Robinson C, et al. Overlapping functions of components of a bacterial Sec-independent protein export pathway. Embo J 1998;17(13):3640–3650.

11. Weiner JH, Bilous PT, Shaw GM, Lubitz SP, Frost L, et al. A novel and ubiquitous system for membrane targeting and secretion of cofactor-containing proteins. Cell 1998;93(1):93–101.

12. Sargent F, Stanley NR, Berks BC, Palmer T. Sec-independent protein translocation in *Escherichia coli*: a distinct and pivotal role for the TatB protein. J Biol Chem 1999;274(51):36073–36082.

13. Bogsch EG, Sargent F, Stanley NR, Berks BC, Robinson C, Palmer T. An essential component of a novel bacterial protein export system with homologues in plastids and mitochondria. J Biol Chem 1998;273(29):18003–18006.

14. Jack RL, Sargent F, Berks BC, Sawers G, Palmer T. Constitutive expression of *Escherichia coli tat* genes indicates an important role for the twin-arginine translocase during aerobic and anaerobic growth. J Bacteriol 2001;183(5):1801–1804.

15. Alcock F, Stansfeld PJ, Basit H, Habersetzer J, Baker MA, et al. Assembling the Tat protein translocase. Elife 2016;5.

16. Rodriguez F, Rouse SL, Tait CE, Harmer J, De Riso A, et al. Structural model for the protein-translocating element of the twin-arginine transport system. Proc Natl Acad Sci U S A 2013;110(12):E1092–1101.

17. Zhang Y, Wang L, Hu Y, Jin C. Solution structure of the TatB component of the twin-arginine translocation system. Biochim Biophys Acta 2014;1838(7):1881–1888.

18. Ramasamy S, Abrol R, Suloway CJ, Clemons WM, Jr. The glove-like structure of the conserved membrane protein TatC provides insight into signal sequence recognition in twin-arginine translocation. Structure 2013;21(5):777–788.

19. Mori H, Cline K. A twin arginine signal peptide and the pH gradient trigger reversible assembly of the thylakoid [Delta]pH/Tat translocase. J Cell Biol 2002;157(2):205–210.

20. Alami M, Luke I, Deitermann S, Eisner G, Koch HG, et al. Differential interactions between a twin-arginine signal peptide and its translocase in *Escherichia coli*. Mol Cell 2003;12(4):937–946.

21. Alcock F, Baker MA, Greene NP, Palmer T, Wallace MI, Berks BC. Live cell imaging shows reversible assembly of the TatA component of the twin-arginine protein transport system. Proc Natl Acad Sci U S A 2013;110(38):E3650–3659.

22. Bolhuis A, Mathers JE, Thomas JD, Barrett CM, Robinson C. TatB and TatC form a functional and structural unit of the twin-arginine translocase from *Escherichia coli*. J Biol Chem 2001;276(23):20213–20219.

23. Leake MC, Greene NP, Godun RM, Granjon T, Buchanan G, et al. Variable stoichiometry of the TatA component of the twin-arginine protein transport system observed by in vivo single-molecule imaging. Proc Natl Acad Sci U S A 2008;105(40):15376–15381.

24. Zoufaly S, Frobel J, Rose P, Flecken T, Maurer C, et al. Mapping precursor-binding site on TatC subunit of twin arginine-specific protein translocase by site-specific photo cross-linking. J Biol Chem 2012;287(16):13430–13441.

25. Blummel AS, Haag LA, Eimer E, Muller M, Frobel J. Initial assembly steps of a translocase for folded proteins. Nat Commun 2015;6:7234.

26. Habersetzer J, Moore K, Cherry J, Buchanan G, Stansfeld PJ, Palmer T. Substrate-triggered position switching of TatA and TatB during Tat transport in *Escherichia coli*. Open Biol 2017;7(8).

27. Gerard F, Cline K. The thylakoid proton gradient promotes an advanced stage of signal peptide binding deep within the Tat pathway receptor complex. J Biol Chem 2007;282(8):5263–5272.

28. Hamsanathan S, Anthonymuthu TS, Bageshwar UK, Musser SM. A Hinged Signal Peptide Hairpin Enables Tat-Dependent Protein Translocation. Biophys J 2017;113(12):2650–2668.

29. Gerard F, Cline K. Efficient twin arginine translocation (Tat) pathway transport of a precursor protein covalently anchored to its initial cpTatC binding site. J Biol Chem 2006;281(10):6130–6135.

30. Panahandeh S, Maurer C, Moser M, DeLisa MP, Muller M. Following the path of a twin-arginine precursor along the TatABC translocase of *Escherichia coli*. J Biol Chem 2008;283(48):33267–33275.

31. Aldridge C, Ma X, Gerard F, Cline K. Substrate-gated docking of pore subunit Tha4 in the TatC cavity initiates Tat translocase assembly. J Cell Biol 2014;205(1):51–65.

32. Dabney-Smith C, Cline K. Clustering of C-terminal stromal domains of Tha4 homo-oligomers during translocation by the Tat protein transport system. Mol Biol Cell 2009;20(7):2060–2069.

33. Dabney-Smith C, Mori H, Cline K. Oligomers of Tha4 organize at the thylakoid Tat translocase during protein transport. J Biol Chem 2006;281(9):5476–5483.

34. Rose P, Frobel J, Graumann PL, Muller M. Substrate-dependent assembly of the Tat translocase as observed in live *Escherichia coli* cells. PLoS One 2013;8(8):e69488.

35. Bruser T, Sanders C. An alternative model of the twin arginine translocation system. Microbiol Res 2003;158(1):7–17.

36. Casadaban MJ, Cohen SN. Lactose genes fused to exogenous promoters in one step using a Mu-lac bacteriophage: in vivo probe for transcriptional control sequences. Proc Natl Acad Sci U S A 1979;76(9):4530–4533.

37. Wexler M, Sargent F, Jack RL, Stanley NR, Bogsch EG, et al. TatD is a cytoplasmic protein with DNase activity. No requirement for TatD family proteins in sec-independent protein export. J Biol Chem 2000;275(22):16717–16722.

38. Lee PA, Orriss GL, Buchanan G, Greene NP, Bond PJ, et al. Cysteine-scanning mutagenesis and disulfide mapping studies of the conserved domain of the twin-arginine translocase TatB component. J Biol Chem 2006;281(45):34072–34085.

39. Liu H, Naismith JH. An efficient one-step site-directed deletion, insertion, single and multiple-site plasmid mutagenesis protocol. BMC Biotechnol 2008;8:91.

40. Engler C, Gruetzner R, Kandzia R, Marillonnet S. Golden gate shuffling: a one-pot DNA shuffling method based on type IIs restriction enzymes. PLoS One 2009;4(5):e5553.

41. Kneuper H, Maldonado B, Jager F, Krehenbrink M, Buchanan G, et al. Molecular dissection of TatC defines critical regions essential for protein transport and a TatB-TatC contact site. Mol Microbiol 2012;85(5):9459–61.

42. Huang Q, Alcock F, Kneuper H, Deme JC, Rollauer SE, et al. A signal sequence suppressor mutant that stabilizes an assembled state of the twin arginine translocase. Proc Natl Acad Sci U S A 2017;114:E1958–E1967.

43. Keller R, de Keyzer J, Driessen AJ, Palmer T. Co-operation between different targeting pathways during integration of a membrane protein. J Cell Biol 2012;199(2):303–315.

44. Cleon F, Habersetzer J, Alcock F, Kneuper H, Stansfeld PJ, et al. The TatC component of the twin-arginine protein translocase functions as an obligate oligomer. Mol Microbiol 2015;98(1):111–129.

45. Dominguez C, Boelens R, Bonvin AM. HADDOCK: a protein-protein docking approach based on biochemical or biophysical information. J Am Chem Soc 2003;125(7):1731–1737.

46. Jo S, Kim T, Iyer VG, Im W. CHARMM-GUI: a web-based graphical user interface for CHARMM. J Comput Chem 2008;29(11):1859–1865.

47. Lindahl E, Abraham MJ, Hess B, van der Spoel D. GROMACS Documentation - Release 2019.2. GROMACS Doc. - Release 2019.2 2–6072019.

48. Huang J, MacKerell AD, Jr. CHARMM36 all-atom additive protein force field: validation based on comparison to NMR data. J Comput Chem 2013;34(25):2135–2145.

49. Bussi G, Donadio D, Parrinello M. Canonical sampling through velocity rescaling. J Chem Phys 2007;126(1):014101.

50. Parrinello M, Rahman A. Polymorphic transitions in single crystals: A new molecular dynamics method. J Applied Phys 1981;52:7182–190.

51. Hess B, Bekker H, Berendsen HJC, Fraaije JGEM. LINCS: A Linear Constraint Solver for Molecular Simulations. J Comp Chem 1997;18:1463–1472.

52. Darden T, York D, Pedersen L. Particle mesh Ewald: An Nlog(N) method for Ewald sums in large systems. J Chem Phys 1993;98:10089–10092.

53. DeLano WL. Pymol: An open-source molecular graphics tool. CCP4 Newsl protein Crystallogr 2002;40:82–92.

54. Ize B, Stanley NR, Buchanan G, Palmer T. Role of the *Escherichia coli* Tat pathway in outer membrane integrity. Mol Microbiol 2003;48(5):1183–1193.

55. Maldonado B, Kneuper H, Buchanan G, Hatzixanthis K, Sargent F, et al. Characterisation of the membrane-extrinsic domain of the TatB component of the twin arginine protein translocase. FEBS Lett 2011;585(3):478–484.

56. Richter S, Bruser T. Targeting of unfolded PhoA to the TAT translocon of *Escherichia coli*. J Biol Chem 2005;280(52):42723–42730.

57. Buchanan G, de Leeuw E, Stanley NR, Wexler M, Berks BC, et al. Functional complexity of the twin-arginine translocase TatC component revealed by site-directed mutagenesis. Mol Microbiol 2002;43(6):1457–1470.

