## Supplemental Tables for "Characterisation of a TatA/TatB binding site on the TatC component of the *Escherichia coli* twin arginine translocase"

| NAME | SEQUENCE | USE* |
|---|---|---|
| ESN15 | CTCGCAAACGCTGTTGGCG**TTC**CCGATGTACTGTCTGTTTGAAATC | QC |
| ESN16 | CTCGCAAACGCTGTTGGCG**TGG**CCGATGTACTGTCTGTTTGAAATC | QC |
| ESN18 | CGCCAACAGCGTTTGCGAGAAGACATCCGGCGGCGTCAGCAAC | QC |
| ESN19 | CTTCTCACGCTTTTACGTTGGTAAAGGGCGAAATCGGGAAGAGG | QC |
| ESN21 | CAACGTAAAAGCGTGAGAAGAAGACACCGAT**AAA**AAACAGACAGTAC | QC |
| ESN22 | CAACGTAAAAGCGTGAGAAGAAGACACCGAT**CCA**AAACAGACAGTAC | QC |
| ESN25 | CTGTTGGCGATCCCGATGTACTGTCTGTTTGAAATCGGTGTCTTCTTC | QC |
| ESN26 | CATCGGGATCGCCAACAG**AAA**TTGCGAGAAGACATCCGGCGGCGTC | QC |
| ESN27 | CATCGGGATCGCCAACAG**CCA**TTGCGAGAAGACATCCGGCGGCGTC | QC |
| ESN33 | GGTGCCTCACTGATTAAGCATTGG | SEQ |
| ESN39 | CTGCATGTGTCAGAGGTTTTCACCGTCATCACCGAAACGCGCGAG | QC |
| ESN40 | GAAAACCTCTGACACATGCAGCTCCCG**T**AGACGGTCACAGCTTGTC | QC |
| ESN41 | CTCGCAAACGCTGTTGGCG**TGG**CCGATGTACTGTCTGTTT**TGG**ATC | QC |
| ESN43 | CTCGCAA**TTT**CTGTTGGCG**TGG**CCGATGTACTGTCTGTTT**TGG**ATC | QC |
| ESN44 | CGCCAACAG**AAA**TTGCGAGAAGACATCCGGCGGCGTCAGCAAC | QC |
| ESN45 | CATATGGTGCACTCTCAGTAC | SEQ |
| ESN47 | CTTGAGCGATGAAAAAATCAGCGTG | CONS |
| ESN48 | GCAACTGGTCTCT*ACCG*CGAGA**T**CCACGCTCACCG | CONS |
| ESN49 | GCAACTGGTCTCT*ATAG*GGAGAC**T**GGAATTCTGTC | CONS |
| ESN50 | GCTGATAAATCTGGAGCCGGTGAG | CONS |
| ESN51 | GCAACTGGTCTCT*GCAT*TCGTTGTCGGGATGTTG | CONS |
| ESN52 | CACGCAGGTCTCT*CGTT*TTCCTCTTCCCGATTTCG | CONS |
| ESN65 | CTCGCAAACGCTGTTGGCGATCCCGATGTAC**NNN**CTGTTTGAAATC | QC |
| ESN74 | GCAACTGGTCTCT*CTAC*CACAGAGGAGGATCCATG | CONS |
| ESN75 | CACGCAGGTCTCA*TCGT*CGACAGACATGCTTAAGG | CONS |
| ESN83 | GTGCAGGTATCCACCGACATC | SEQ |
| ESN91 | CATCGGGATCGCCAACAG**TTT**TTGCGAGAAGACATCCGGCGGCGTC | QC |
| ESN93 | CTCGCAAACGCTGTTGGCG**CGT**CCGATGTAC**GCT**CTGTTTGAAATC | QC |
| ESN104 | AGTTAGCTCACTCATTAGGC | SEQ |
| ESN109 | CTCGCAAACGCTGTTGGCGATCCCGATGTACTGT**CAG**TTTGAAATC | QC |
| ESN112 | CCGTATGTGCTGGTTGGT*GCAT*TGAGACCGAGCGCAACGCAATTAATGTG | CONS |
| ESN113 | CGCTTTCTGCTTCAGCGT*CGTT*AGAGACCTTAGCGCCATTCGCCATTCAG | CONS |
| ESN114 | AACGACGCTGAAGCAGAAAGCG | CONS |
| ESN115 | ATGCACCAACCAGCACATACGG | CONS |
| ESN118 | AAGAACATCGATTTTCCATGGCAG | CONS |
| ESN119 | CGACTCACTATAGGGAGAGCGGC | CONS |
| ESN123 | CGGCTTTGTTTAATCATCAT*CTAC*CGAGACCGAGCGCAACGCAATTAATGTG | CONS |
| ESN124 | GTGATAAGCGGTTGAGTA*TCGT*CGAGACCTTAGCGCCATTCGCCATTCAG | CONS |
| ESN125 | GGTAGATGATGATTAAACAAAGCCG | CONS |
| ESN126 | GACGATACTCAACCGCTTATCAC | CONS |
| ESN128 | GAGCGCAACGCAATTAATGTG | CONS |
| ESN129 | TTAGCGCCATTCGCCATTCAG | CONS |
| ESN132 | GTTACACGCAGGTCTCT*CGTT*TTCCTCTTCCCGATTTCG | CONS |
| ESN139 | CTGAATGGCGAATGGCGCTAAGGTCTCT*AACG*ACGCTGAAGCAGAAAGCG | CONS |
| ESN140 | GATGTCGGTGGATACCTGCAC | SEQ |
| ESN152 | CACATTAATTGCGTTGCGCTCGGTCTCT*TTGC*GAGAAGACATCCGGCGGCGTC | CONS |
| ESN153 | CTGAATGGCGAATGGCGCTAAGGTCTCG*CTTT*TACGTTGGTAAAGGGCGAAATC | CONS |
| ESN154 | CTCGCAA**TTT**CTGTTGGCG**TGG**CCGATGTAC**GCT**CTGTTT**TGG**ATC | QC |
| ESN160 | CTCGCAAACGCTGTTGGCGATC**CGT**ATGTAC**GCT**CTGTTTGAAATC | QC |
| ESN161 | CTCGCAAACG**CGGCTC**GCGATCCCGATGTAC**GCT**CTGTTTGAAATC | QC |
| ESN162 | CGC**GAGCCG**CGTTTGCGAGAAGACATCCGGCGGCGTCAGCAAC | QC |
| ESN163 | CGC**AAA**CAGCGTTTGCGAGAAGACATCCGGCGGCGTCAGCAAC | QC |
| ESN164 | CTCGCAAACGCTG**TTT**GCGATCCCGATGTAC**GCT**CTGTTTGAAATC | QC |
| ESN167 | CTCGCAAACG**CCGCTC**GCGATCCCGATGTAC**GCT**CTGTTTGAAATC | QC |
| ESN168 | CGC**GAGCGG**CGTTTGCGAGAAGACATCCGGCGGCGTCAGCAAC | QC |
| ESN169 | CTCGCAAACGCTGTTGGCG**AAC**CCGATGTAC**GCT**CTGTTTGAAATC | QC |
| ESN170 | CTCGCAAACGCTGTTGGCGATCCCG**CTG**TAC**GCT**CTGTTTGAAATC | QC |

| | | |
|---|---|---|
| ESN174 | CACATTAATTGCGTTGCGCTCGGTCTCG*TCAG*CAACATCCCGACAACGAATGC | CONS |
| ESN175 | CACATTAATTGCGTTGCGCTCGGTCTCG*TCAG*CAA**GCA**CCCGACAACGAATGC | CONS |
| ESN176 | ATGGCGCTAAGGTCTCG*CTGA*CGCCGCCGGATGTCTTCTCGCAA | CONS |
| ESN177 | ATGGCGCTAAGGTCTCG*CTGA*CGCCGCCGGATGTCTGCTCGCAA | CONS |
| ESN178 | GCATTCGTTGTCGGGATGTTGCTGAC | CONS |
| ESN180 | GTCAGCAACATCCCGACAACGAATGC | CONS |
| ESN182 | CGCTTTCTGCTTCAGCGTCGTT | CONS |
| ESN187 | CTCGCAAACGCTGTTGGCGATCCCGATGTAC**GCTCCG**TTTGAAATC | QC |
| ESN189 | CTCGCAAACGCTGTTGGCGATCCCGATGTAC**GCT**CTGTTTGAAATC | QC |
| ESN191 | CAACGTAAAAGCGTGAGAAGAAGACACCGAT**ACC**AAACAG**AGC**GTAC | QC |
| ESN192 | CAACGTAAAAGCGTGAGAAGAA**GTC**ACCGATTTCAAACAG**AGC**GTAC | QC |
| ESN199 | CTCGCAAACGCTGTTGGCGATC**CGT**ATGTACTGTCTGTTTGAAATC | QC |
| ESN200 | **CTT**CAACAGCGTTTGCGAGAAGACATCCGGCGGCGTCAGCAAC | QC |
| ESN201 | CTCGCAAACGCTGTTG**AAG**ATCCCGATGTACTGTCTGTTTGAAATC | QC |
| ESN203 | **GCG**CAACAGCGTTTGCGAGAAGACATCCGGCGGCGTCAGCAAC | QC |
| ESN204 | CTCGCAAACGCTGTTG**CGC**ATCCCGATGTACTGTCTGTTTGAAATC | QC |
| ESN207 | CAACGTAAAAGCGTGAGAAGAAGACACCGATTTC**ATC**CAG**AGC**GTAC | QC |
| ESN209 | C**AAA**TCACGCTTTTACGTTGGTAAAGGGCGAAATCGGGAAGAGG | QC |
| ESN210B | CAACGTAAAAGCGTGA**TTT**GAAGACACCGATTTCAAACAG**AGC**GTAC | QC |
| ESN211 | C**GAA**TCACGCTTTTACGTTGGTAAAGGGCGAAATCGGGAAGAGG | QC |
| ESN212 | CAACGTAAAAGCGTGA**TTC**GAAGACACCGATTTCAAACAG**AGC**GTAC | QC |
| ESN213 | CTCGCAAACGCTGTTGGCGATCCCG**CGT**TACTGTCTGTTTGAAATC | QC |
| ESN217 | CATCGGGATCGCCAACAG**TTC**TTGCGAGAAGACATCCGGCGGCGTC | QC |
| ESN218 | CATCGGGATCGCCAACAG**CAG**TTGCGAGAAGACATCCGGCGGCGTC | QC |
| ESN219 | CAACGTAAAAGCGTGAGAAGAAGAC**GTC**GATTTCAAACAGACAGTAC | QC |
| ESN220 | CTCGCAAACGCTG**GAG**GCGATCCCGATGTACTGTCTGTTTGAAATC | QC |
| ESN221 | CGC**CTC**CAGCGTTTGCGAGAAGACATCCGGCGGCGTCAGCAAC | QC |
| ESN225 | CTCGCAAACGCTGTTGGCGATCCCG**CCG**TACTGTCTGTTTGAAATC | QC |
| AL1 | CTCGCAAACGCTGTTGGCGATCCCGATG**TCC**TGTCTGTTTGAAATC | QC |
| AL2 | CAACGTAAAAGCGTGAGAAGAAGACACCGATTTC**GCG**CAGACAGTAC | QC |

**Table S1. Oligonucleotides used in this study**

Note that the term “CONS” is used for plasmid construction by ligation or an assembly technique, i.e. pJET1.2/blunt cloning, NEBuilder, Golden Gate (in the latter case, *Bsa*I recognition sites are underlined and the corresponding quadruplets are in italics); “SEQ”: used for sequencing (ESN33 and ESN45 are specific to pTAT1d; ESN104 to pTAT101; ESN83 and ESN140 are internal to *tatC*); “QC”: used for QuickChange (see Table 1). In all primers, mutated bases or codons are highlighted in bold.

| TatC stop codon substitutions |
|---|
| L206stop (isolated 14 times) |
| S214stop (isolated 3 times) |
| Q215stop |
| P210L, Q215stop |
| L218stop (isolated 20 times) |
| V203D, L218stop |
| L206M, L218stop |
| T208A, L218stop |
| L44R, L218stop |
| Y223stop |
| V202D, Y223stop |
| M222I, Y223stop |
| C224stop – (isolated twice) |
| E227stop |
| M205K, E227stop |
| F226V, E227stop |
| F213S, I228N, Y236stop |
| M205K, P221T, K239stop |
| TatC frameshifts |
| F213::frameshift |
| Q215::frameshift |
| T216::frameshift |
| S214P, T216::frameshift |
| L217::frameshift (isolated 3 times) |
| L218::frameshift (isolated twice) |
| A219::frameshift |
| T208A, Q215H, A219::frameshift |
| I220::frameshift (isolated twice) |
| P221::frameshift |
| M222::frameshift |
| L217M, C224::frameshift |
| F226::frameshift (isolated 3 times) |
| L207R, I228::frameshift |
| T216R, V230::frameshift |
| V203A,T208A, P209Q, C224R, K239::frameshift |
| TatC in-frame deletion |
| Δ(amino acids 227-234) |

**Table S2. Stop codon, frame-shift and in-frame deletion mutations isolated from error prone and scanning mutagenesis library screens for inactivating substitutions in TatC**

**TMH6.** All stop codon mutations were isolated from the error-prone PCR library screen, whereas the in-frame deletion was isolated from the scanning mutagenesis library. Frame-shift mutations were isolated from both libraries.
